# A novel monocarboxylate transporter involved in 3-hydroxykynurenine transport for ommochrome coloration

**DOI:** 10.1101/2023.06.01.543243

**Authors:** Hirosumi Uchiyama, Yoko Takasu, Minoru Moriyama, Kazutoshi Yoshitake, Hironobu Uchiyama, Tetsuya Iizuka, Keiro Uchino, Genta Okude, Yutaka Banno, Seigo Kuwazaki, Kimiko Yamamoto, Shunsuke Yajima, Hideki Sezutsu, Toshiki Tamura, Ryo Futahashi, Mizuko Osanai-Futahashi

## Abstract

Ommochromes are widespread pigments in invertebrates utilized for screening pigments in compound eyes and for reddish coloration in epidermis and wings. Ommochromes are derived from 3-hydroxykynurenine (3OHK), which is incorporated into cells from hemolymph or synthesized from tryptophan within cells. While the synthetic pathway from tryptophan to 3OHK has been well characterized, the gene responsible for cellular uptake of 3OHK has been poorly understood. In the silkworm *Bombyx mori*, adult compound eyes and eggs contain a mixture of ommochrome pigments. By using positional cloning method, we found that a novel monocarboxylate transporter, 3-hydroxykynurenine transporter (3OHKT), is responsible for the recessive mutant *maternal brown of Tsujita* (*b-t*) of *B. mori*. In *b-t* mutant, the color of the eggs is light brown, whereas the color of the compound eyes is normal, and we identified a 2-kb deletion in *3OHKT* gene. TALEN-mediated knockout of *3OHKT* gene produced the same coloration phenotype as *b-t* mutant, and the complementation test between *b-t* mutant and *3OHKT* knockout strain proved that *3OHKT* is responsible for *b-t* phenotype. 3OHKT protein was localized in the cellular membrane, and LC-MS analysis indicated that the uptake of 3OHK from hemolymph into the ovary was suppressed in the *b-t* mutant. Moreover, we confirmed that *3OHKT* gene is specifically expressed at the reddish region and the time of pigmentation in the pupal wing of nymphalid butterflies. RNA interference of *3OHKT* prevented reddish pigmentation in wings, highlighting its general involvement in ommochrome-based pigmentation other than compound eyes.

**Significance:** Ommochromes are widely distributed pigments in invertebrates and are synthesized from intracellular tryptophan or 3-hydroxykynurenine (3OHK). Ommochrome-based red markings on butterfly wings are often used for sexual selection, warning colors and mimicry. Most genes involved in the ommochrome synthesis pathway have been elucidated from analyses of eye color mutants in *Drosophila*. However, this study reveals that the ommochrome synthesis pathway has a different genetic repertoire depending on the tissues, and that the novel monocarboxylate transporter identified in this study has a major role in ommochrome pigmentation other than in compound eyes. In particular, our results suggest that classical ommochrome-related genes are rarely involved in the wing pigmentation of the nymphalid butterflies.

## Introduction

Ommochromes represent a major pigment group in invertebrates, comprising yellow, orange, red, and purple pigments (Figon and Casas, 2019). They are distributed in compound eyes as screening pigments in various insects, and often found in the epidermis and wings (Linzen, 1974; Figon and Casas, 2019; Williams et al., 2019; Futahashi and Osanai-Futahashi, 2021). In Nymphalidae, the largest family of butterflies, ommochromes are often detected in red and/or orange markings of wings (Fig. 1A, B), which function as sexual selection, warning colors, and mimicry (Nijhout, 1991). The genes involved in ommochrome biosynthesis have been identified primarily by genetic analyses of eye color mutants in the fruit fly *Drosophila melanogaster*. Three enzymes, encoded by *vermilion, kynurenine formamidase*, and *cinnabar*, convert the ommochrome precursor tryptophan to 3-hydroxykynurenine (3OHK) within cells (Searles et al., 1990; Searles and Voelker, 1986; Warren et al., 1996). Subsequently, 3OHK is incorporated into pigment granules namely ommochromasomes by the heterodimeric ABC transporters encoded by *scarlet* and *white* (Pepling and Mount, 1990; Tearle et al., 1989), and converted into final pigments by heme peroxidase encoded by *cardinal* within ommochromasomes (Harris et al., 2011). This pathway is conserved in other insects, such as the red flour beetle *Tribolium castaneum* (Broehan et al., 2013; Grubbs et al., 2015; Lorenzen et al., 2002), the Western tarnished plant bug, *Lygus hesperus* (Brent and Hull, 2019), and the silkworm *Bombyx mori* (Kômoto et al., 2009; Quan et al., 2007; Tatematsu et al., 2011). Transplantation and tracer experiments in several insects have shown that ommochromes in some tissues (e.g., ovaries, wings) can be synthesized by the uptake of not only tryptophan but also 3OHK from hemolymph (Sonobe and Ohnishi, 1970; Linzen, 1974; Koch, 1991; Koch, 1993). In the nymphalid butterfly *Araschnia levana*, 3OHK is incorporated more efficiently than tryptophan in the red regions of pupal wings (Koch, 1993). Although the synthetic pathway from tryptophan to 3OHK has been well characterized, the gene responsible for the cellular uptake of 3OHK has been poorly understood.

**Fig. 1.**
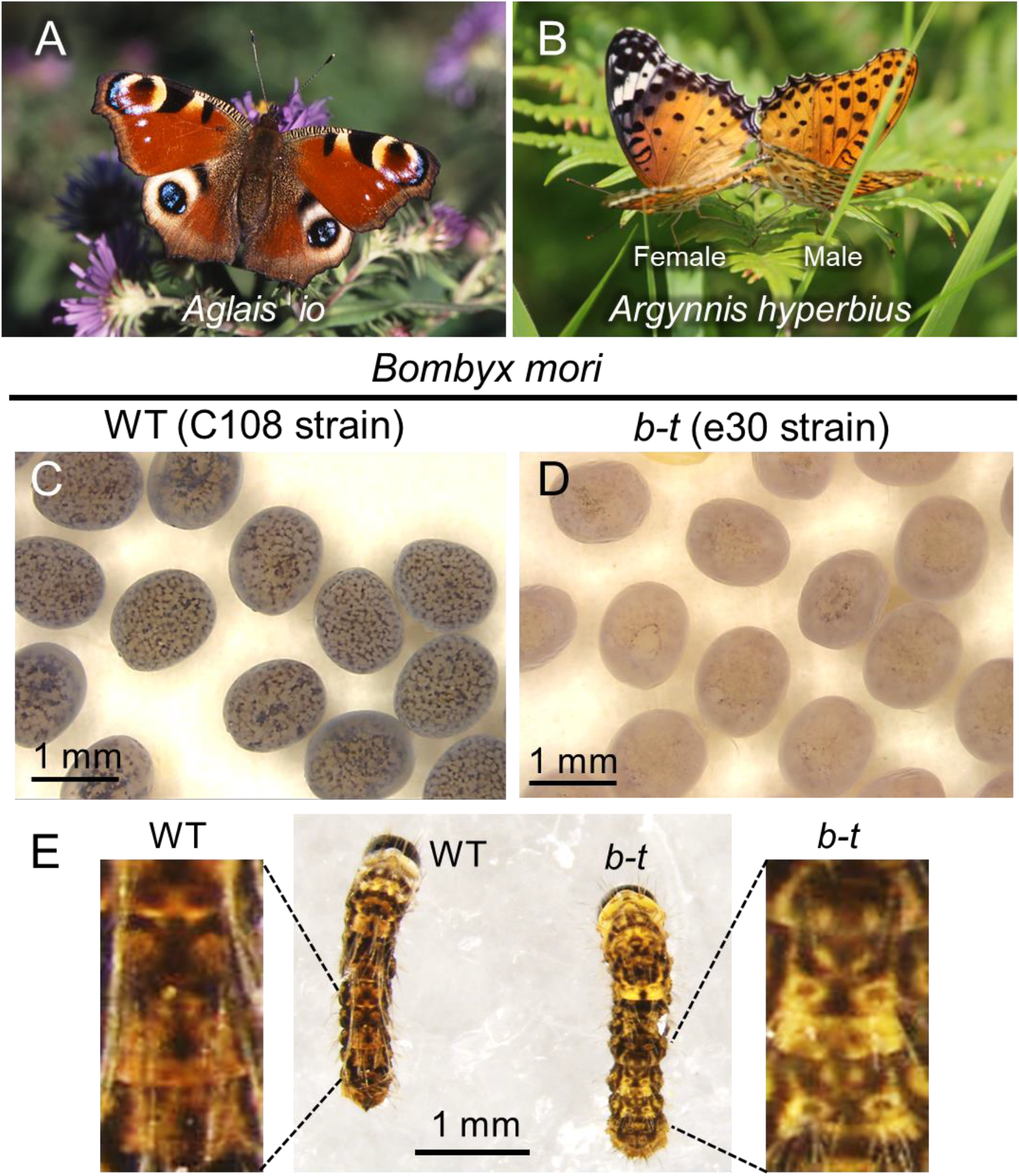
Examples of ommochrome-based coloration in butterflies and moths. (A) The peacock butterfly *Aglais io*. (B) The Indian fritillary butterfly *Argynnis hyperbius* with prominent sexual dimorphism in wing coloration. (C) Eggs of wild-type (WT) of the silkworm *Bombyx mori*. (D) Eggs of *b-t* mutant strain of *B. mori*. (E) The 1st instar larva of wild-type (WT) of *B. mori*. (F) The 1st instar larva of *b-t* mutant strain of *B. mori*.

While only ommatin-type ommochrome pigments have been identified from compound eyes of *Drosophila, B. mori* has multiple types of ommochrome pigments which give their eggs and compound eyes dark coloration. Genes responsible for several mutants of ommochrome biosynthesis in *B. mori* have been identified (Quan et al., 2007; Komoto et al., 2009; Tatematsu et al., 2011; Osanai-Futahashi et al., 2016; Tomihara et al., 2021), and the *red egg* gene important for ommin-type ommochrome biosynthesis does not have homologs in *Drosophila* (Osanai-Futahashi et al., 2012). However, the gene responsible for the *B. mori* mutant *maternal brown of Tsujita* (*b-t*), whose egg color is light brown instead of a normal dark coloration, has not been identified.

*b-t* is a recessive mutant whose lighter egg color depends on the mother’s genotype, and the *b-t* mutation has been mapped to genetic linkage group 13 (chromosome 13) (Doira et al., 1981). In this study, based on positional cloning, TALEN-mediated gene knockout, and complementation tests, we identify that the gene responsible for *b-t* is novel monocarboxylate transporter, designated as *3-hydroxykynurenine transporter* (*3OHKT*). Moreover, we confirmed that *3OHKT* gene is specifically expressed in the red region of the wings of nymphalid butterflies and RNA interference (RNAi) of *3OHKT* inhibits reddish wing pigmentation, indicating that this gene is widely involved in coloration by ommochrome pigments.

## Results

### *b-t* mutations reduce the ommochrome-based pigmentation in *B. mori* except for the compound eyes

*b-t* mutant has pale brown eggs in contrast to dark purple-colored eggs of the wild-type (Fig. 1C, D), while the color of the adult compound eyes is indistinguishable between wild-type and *b-t* as previously reported (*SI Appendix*, Fig. S1A) (Doira et al., 1981). In *Bombyx mori*, ommochrome pigments also accumulate in the epidermis of the 1^st^ instar larva, and the larval ganglion (Osanai-Futahashi et al., 2012; Tomihara et al., 2021). We found that ommochrome-based pigmentation in epidermis of the 1st instar larvae and the larval ganglion was lighter in *b-t* strain than in wild-type (Fig. 1E; *SI Appendix*, Fig. S1B). The coloration of the neonatal feces was also lighter in *b-t* than in wild-type (*SI Appendix*, Fig. S1C). The color of the neonatal feces likely reflects that the egg pigment in the serosa is swallowed by the larva about two days before hatching.

### Positional cloning of the *b-t* locus in *B. mori*

In order to identify the genomic region linked to *b-t*, we performed double digest restriction-site associated DNA sequencing (ddRAD-seq). We obtained F1 heterozygous males from a cross between a single female of *b-t* mutant strain (e30) and a single male of wild-type (C108 strain), and backcrossed the F1 heterozygous males with *b-t* mutant females, and subjected BC1, F1, parental individuals for dd-RADseq analyses. We narrowed down the *b-t* linked region to within 523 kbp on chromosome 13 containing nine predicted genes (Fig. 2A, B). To determine the candidate genes, we conducted RNA sequencing analysis of eggs at 0, 24, 48 and 72 after oviposition using *b-t* and wild-type (p50T and C108 strains), and compared the sequence, expression, and splicing pattern of 9 candidate genes among *b-t*-linked region (Fig. 2C; *SI Appendix*, Fig. S2). Among these genes, only a novel monocarboxylate transporter gene (*KWMTBOMO07980*) displayed a difference in splicing pattern between *b-t* and wild-type strains (Fig. 3A). In *b-t* strain, the region downstream of the original exon 9 was used as exon 9, causing alteration of the amino acid sequence of the C-terminus (Fig. 3A). This gene was expressed prior to other ommochrome related genes (e.g., *cardinal, red egg*, and *scarlet*) during egg pigmentation in wild-type strains (Fig. 2C). By genomic PCR, we found that 2 kb region which includes a part of intron 8 and most of the exon 9 of wild-type, was deleted in *b-t* strain (Fig. 3B). The wild-type KWMTBOMO07980 protein, hereafter designated as 3-hydroxykynurenine transporter (3OHKT), was predicted to be a 12-pass transmembrane protein, whereas *b-t* type protein was predicted to have only 11 transmembrane domains (*SI Appendix*, Fig. S3). 3OHKT protein belongs to a monocarboxylate transporter family, and homologs of *B. mori* 3OHKT were found across insect species (Fig. 3C; *SI Appendix*, Fig. S4). *B. mori* had eight closely related genes including *3OHKT*, four of which were located in tandem on chromosome 13 (Fig. 3D). The other 4 genes were located on chromosome 5, one of which encoded a putative uric acid transporter responsible for the silkworm oily mutant *oc* (Wang et al., 2020).

**Fig. 2.**
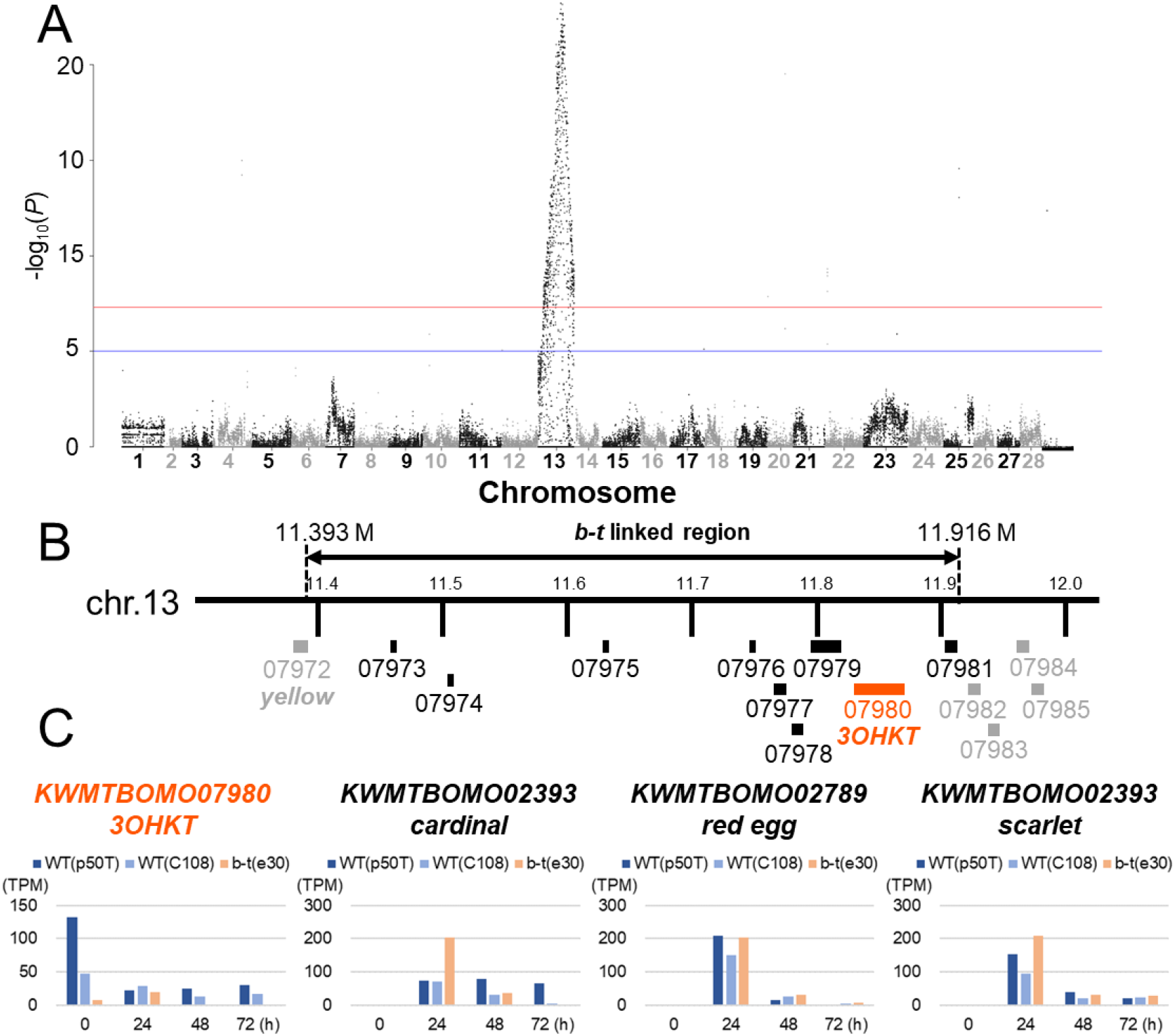
Positioinal cloning of *b-t* candidate region in the silkworm *B. mori*. (A) Manhattan plot for ddRAD-seq analysis in F2 individuals. The blue line and red line indicate p-value = 1.0 × 10^−5^ and 5.0 × 10^−8^, respectively. A significant (p-value < 5.0 × 10^−8^) peak was observed only on chromosome 13. (B) A physical map of the *b-t*-linked region. The *b-t*-linked region was narrowed within the 523 kb region containing 9 predicted genes (e.g. 07980: *KWMTBOMO07980*). (C) Expression analysis of *KWMTBOMO07980* and 3 ommochrome-related genes in embryos of *B. mori*. The number on the x axis indicate hours after oviposition.

**Fig. 3.**
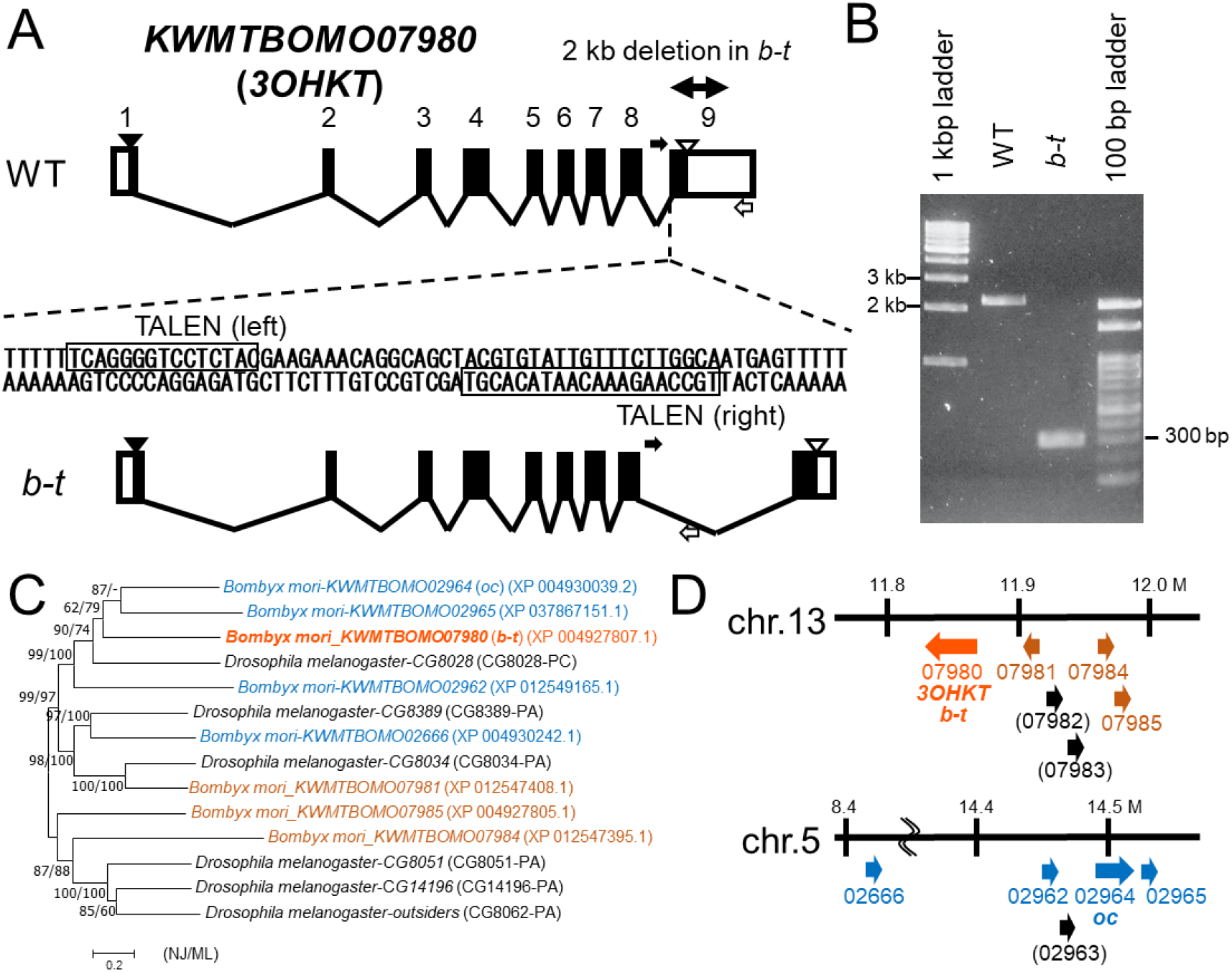
KWMTBOMO07980 exon-intron structure and genomic deletion of the b-t strain in B. mori. (A) Exon-intron structure of *KWMTBOMO07980* (*3OHKT*) gene of wild-type (p50T strain, top) and *b-t* (bottom), and the TALEN binding sites, based on embryonic RNA-seq analysis. The open reading frame, untranslated regions, introns, start codons, and stop codons are indicated by solid boxes, open boxes, diagonal lines, black arrowheads and open arrowheads, respectively. The primers used for genomic PCR in (B) are indicated by solid and open arrows. (B) Genomic PCR of *KWMTBOMO07980* of wild-type (p50T) and *b-t* using primers b-t_intron F7 (black arrow), and b-t_exon9 R2 (open arrow). 2 kb deletion in *b-t* was identified. (C) Phylogenetic relationship of *KWMTBOMO07980* (*3OHKT*) gene homolog in *B. mori* and the fruit fly *Drosophila melanogaster*. A neighbor-joining phylogeny is shown. On each node, bootstrap values are indicated in the order of neighbor-joining method/maximum-likelihood method. A hyphen indicates value less than 50%. Accession numbers or annotation IDs are in parenthesis. (D) Genomic location of the *B. mori KWMTBOMO07980* homolog genes. Numbers below arrows indicate the numbers of predicted genes (e.g. 07980: *KWMTBOMO07980*). Numbers in parentheses are genes considered to be transposable elements.

### Knockout of *3OHKT* gene by TALEN induces *b-t* phenotype

To investigate whether *KWMTBOMO07980* (*3OHKT*) was the responsible gene for *b-t*, we generated knockout individuals for *3OHKT* gene by transcription activator-like effector nuclease (TALEN). TALEN target sites were designed in exon 9 of wild-type *3OHKT* (Fig. 3A) to disrupt exon 9 like *b-t* mutant. We obtained 78 G1 egg batches from G0 sibling pairing. Because the pale brown egg color of *b-t* mutant is subject to the maternal genotype, the color of the eggs laid by G1 females was examined. Of the 32 G1 females, 18 females laid light brown *b-t*-like eggs (*SI Appendix*, Fig S5). In order to obtain homozygous strain of the knockout allele, G2 individuals obtained from sib-mating of G1 individuals were outcrossed with wild-type (C108), and their F1 offsprings were sib-mated. To select *3OHKT* knockout homozygotes, first instar larvae with pale-colored abdomen were selected from F2 broods which both parents have the same genotype. As a result, *3OHKT* Δ10 (= 10-bp deletion) homozygous strain was established, which laid the pale brown colored eggs similar to that of *b-t* (Fig 4).

**Fig. 4.**
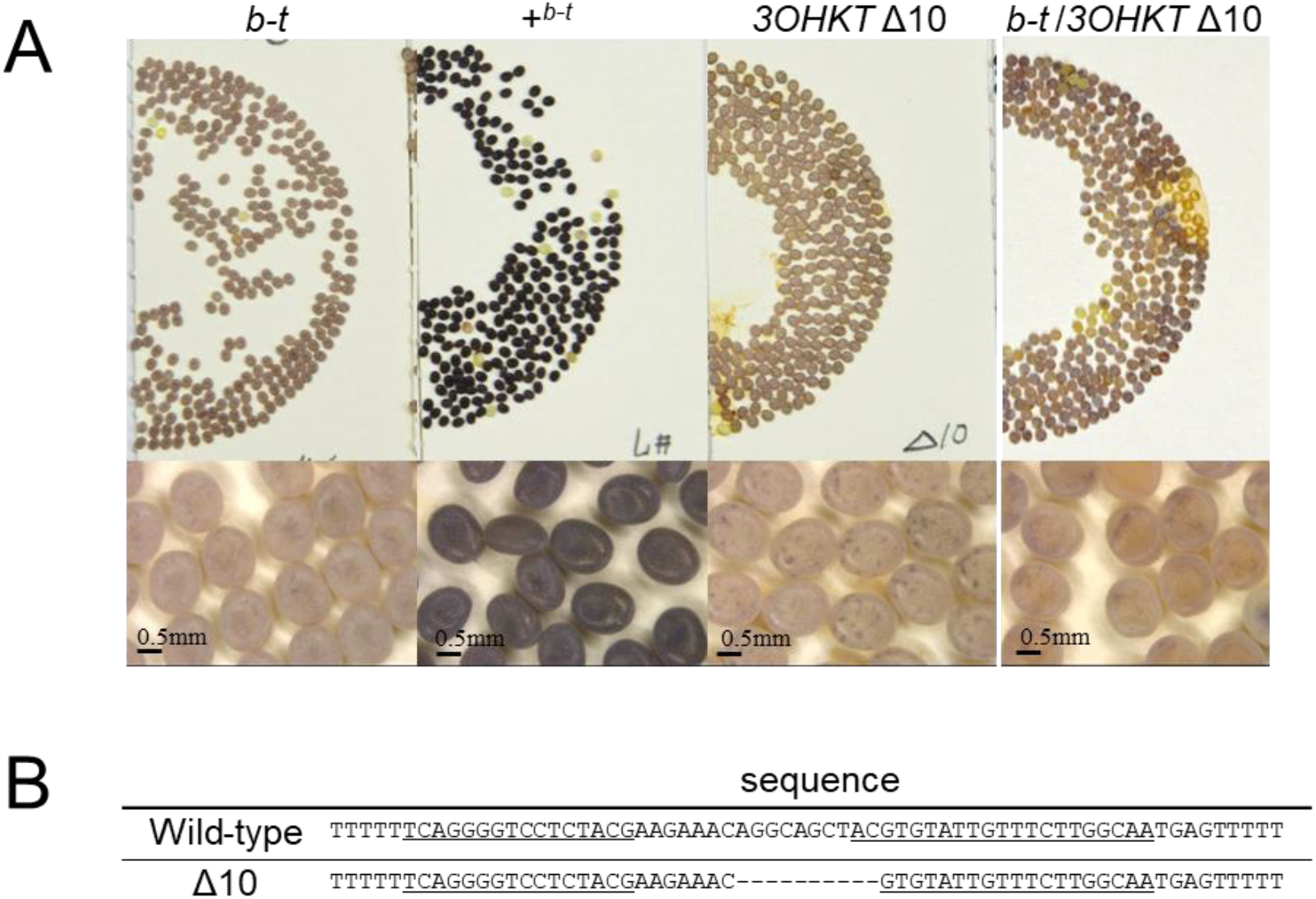
Disruption of the *KWMTBOMO07980* (*3OHKT*) gene using designed TALENs in *B. mori*. (A) Eggs laid by *b-t*, wild-type (C108), *3OHKT* Δ10 homozygous, and *b-t*/*3OHKT* Δ10 transheterozygous. (B) *3OHKT* sequence of wild-type and Δ10 homozygous. TALEN binding sequences are underlined.

To prove that *3OHKT* gene is responsible for the *b-t* phenotype, we performed complementation test by crossing *b-t* mutant females with *3OHKT* knockout males (Fig 4; *SI Appendix*, Fig. S6). The next generation was dissected for inspection of ganglia pigmentation at the larval stage, or sib-mated. The ganglia of the fifth instar larvae were lighter in color than the wild-type like *b-t* (*SI Appendix*, Fig. S6B). The color of eggs obtained from sib-mating was all pale brown like *b-t* (Fig 4; *SI Appendix*, Fig. S6C). The above results demonstrate that the *3OHKT* is responsible for the *b-t* phenotype.

### 3OHKT is involved in 3OHK incorporation into eggs

To examine 3OHKT protein localization, we analyzed the subcellular distribution of the protein by fusing it to a 3xFLAG tag at the amino terminus and expressed in *B. mori* cultured cell BmN4-SID1. The 3xFLAG fusion protein was then visualized by immunofluorescence analysis and confocal microscopy. As a result, 3xFLAG-3OHKT localized on the cell membrane and on vesicle-like structures in the cytoplasm (Fig. 5A).

**Fig. 5.**
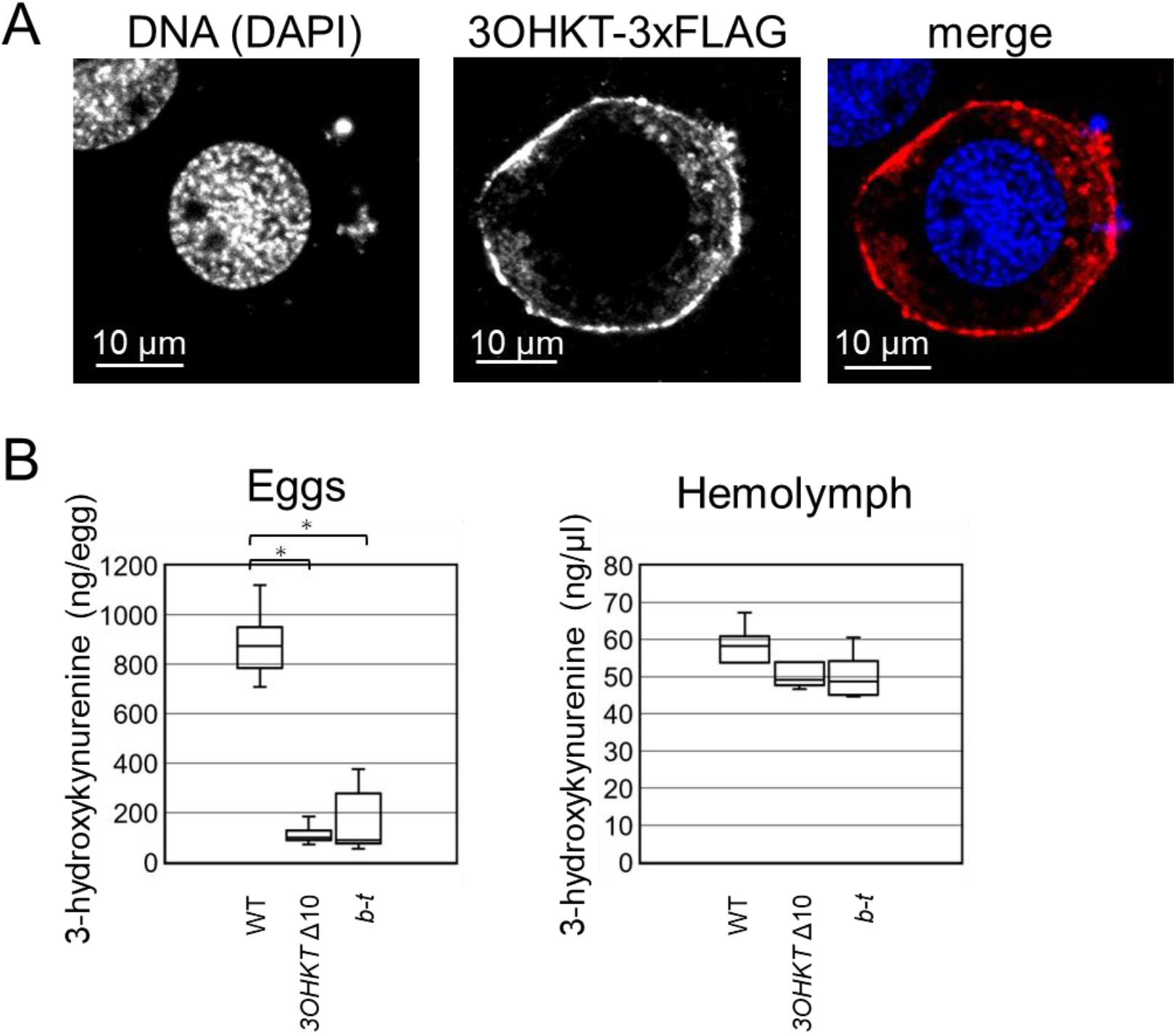
Cellular localization of 3OHKT protein and quantification of 3-hydroxykynurenine in eggs and hemolymph of *B. mori*. (A). Immunocytochemistry of BmN4-SID1 cells expressing 3OHKT-3xFLAG. In the merged image, 3OHKT and DNA is indicated by red and blue, respectively. 3OHKT is localized mainly in the plasma membrane. (B) Quantification of 3-hydroxykynurenine (3OHK) of egg and hemolymph samples. 3OHK was quantified from ten eggs for each batch, and average amount per egg was shown. Replicates of egg samples analyzed for *3OHKT* knockout (Δ10) (left), wild-type (C108) (middle) and *b-t* (right) were 8, 7, and 5, respectively. The number of hemolymph samples for *3OHKT* knockout (Δ10) (left), wild-type (C108) (middle) and *b-t* (right) were 4, 5, and 4, respectively. Error bars indicate SEM. * *p* < 0.05 for Student’s *t*-test.

*B. mori* ovaries have the activity to convert kynurenine to 3OHK, but also incorporate 3OHK from the hemolymph during the pupal stage, when the ovary rapidly develops (Sonobe and Ohnishi, 1970; Linzen, 1974). It has been presumed that the amount of 3OHK in eggs is reduced in the *b-t* strain (Doira et al., 1981). We measured the amount of 3OHK in eggs within 24 hours after oviposition which is before the maternally-derived 3OHK is used up for ommochrome synthesis, and in hemolymph from the female day 4 pupae, which is shortly after 3OHK uptake from hemolymph into ovaries has begun. The amount of 3OHK in wild-type eggs was over sevenfold and fourfold of that in the *3OHKT*Δ10 eggs and *b-t* eggs, respectively (Fig 5B). On the other hand, the amount of 3OHK in the pupal hemolymph did not significantly differ between the three strains (Fig 5B). These results indicate that 3OHKT protein functions mainly in the uptake of 3OHK from hemolymph into eggs.

### *3OHKT* defect affects ommochrome synthesis in eggs but not in the compound eyes

The color of the eggs, the larval ganglion, and the epidermis of *b-t* strain and *3OHKT* knockout mutants was lighter than that of the wild-type. We extracted ommochrome pigments with acidic methanol (HCl/MeOH) from eggs of wild-type strain and *b-t* mutant strains. The color of egg extracts of *b-t* and *3OHKT* Δ10 were markedly lighter than that of the wild type (*SI Appendix*, Fig. S7A-C, G-I). Since ommochrome pigments can change color depending on the redox status (Linzen, 1974), we measured the absorbance spectrum of the pigments using a spectrophotometer with or without an oxidizing (NaNO_2_) or a reducing (ascorbic acid) agent to observe redox-dependent color changes. The absorbance spectra of *3OHKT* Δ10 and *b-t* egg extracts were significantly lower than those of the wild-type strain at wavelength 360, 380, 450, 480, and 530 nm, regardless of addition of oxidizing or reducing agent (*SI Appendix*, Fig. S7D-F, J-L). This suggests that the total amount of ommochrome pigments in eggs is smaller in both *b-t* and *3OHKT* Δ10 compared to wild-type. In contrast, the color of the eye extracts was indistinguishable between wild-type, *3OHKT* Δ10 and *b-t*, and no significant difference was detected in the absorbance spectrum (*SI Appendix*, Fig. S8). We also investigated the expression of *3OHKT* gene in the ovary and compound eye of pupa on the day 4 and day 8. The expression in compound eyes was significantly lower than that in ovaries (*SI Appendix*, Fig S9), which is consistent with the lower contribution of 3OHKT protein to eye pigmentation.

### *3OHKT* is specifically expressed at the stage and region of ommochrome coloration in the wings of the nymphalid butterflies

Next we focused on the ommochrome-related reddish wing coloration of nymphalid butterflies. We performed RNA sequencing using samples of wings and compound eyes during larvae and pupae of the peacock butterfly *Aglais io* (also known as *Nymphalis io* and *Inachis io*) which has reddish-brown wing coloration due to ommochrome pigments ommatin D and rhodommatin (Linzen, 1974). Besides the reddish-brown ommochrome pigmented region, *A. io* has a large eye spot on each wing, and melanic markings on the forewing (Figs. 1A and 6). The ommochrome and melanin pigmentation occurs pupa day 6 and day 7, respectively (Fig. 6A). *3OHKT* gene was preferentially expressed in the red-destined wing regions of the *A. io* pupa at day 6 and day 7, respectively (Fig. 6B), strongly suggesting that *3OHKT* gene is involved in the reddish ommochrome pigmentation of *A. io* wings. In addition to *3OHKT*, genes for the synthesis of 3OHK from tryptophan (e.g., *vermilion, Kynurenine formamidase, cinnabar*) were also expressed in the wings (Fig. 6B; *SI Appendix*, Fig. S10). Meanwhile, genes involved in ommochrome-related genes downstream of 3OHK synthesis, such as *cardinal, red egg*, and *scarlet*, were highly expressed in the compound eyes prior to eye pigmentation, but were hardly expressed in the wings (*SI Appendix*, Fig. S10).

**Fig. 6.**
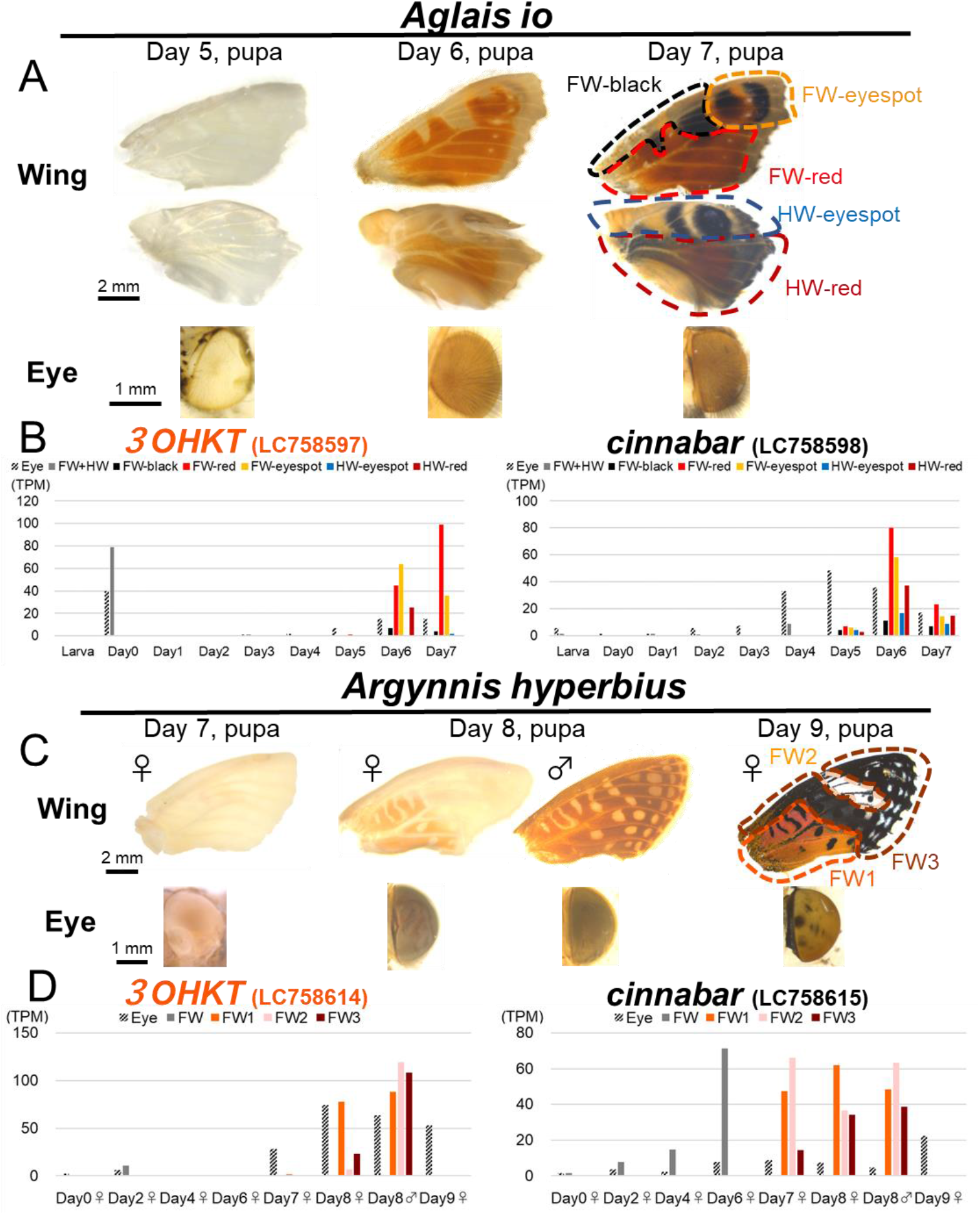
Expression profiles of *3OHKT* and *cinnabar* genes in wings and eyes of peacock butterflies *Aglais io* and *Argynnis hyperbius*. (A) Coloration of wings and compound eyes on days 5-7 after pupation of *A. io*. After day 5, pupal wings were separated for markings as indicated. (B) Expression profiles of *3OHKT* and *cinnabar* in *A. io* (C) Coloration of wings and compound eyes on days 7-9 after pupation of *A. hyperbius*. After day 7, pupal forewings were separated for markings as indicated. In males, the regions corresponding to the female pattern were separated. (D) Expression profiles of *3OHKT* and *cinnabar* in *A. hyperbius*. FW, forewing; HW, hindwing.

We also performed RNA sequencing using samples of pupal wings and compound eyes of the Indian fritillary butterfly *Argynnis hyperbius*, which has orange and black wing coloration with remarkable sexual dimorphism (Figs. 1B and 6). Like *A. io*, ommatin D and rhodommatin are detected in *Argynnis* wings (Linzen, 1974). In wings, *3OHKT* was transiently expressed upon orange pigmentation stage (Day 8, pupa) in the sex-specific orange regions (FW1 for females and FW1, FW2 and FW3 for males) (Fig. 6C, D). It should be noted that *cinnabar* is also expressed in wings, but no significant difference in the expression level was observed during orange pigmentation stage between male and female. Similar to *A. io*, many of the ommochrome-related genes (e.g., *cardinal, red egg, scarlet*) were highly expressed in the compound eyes but were hardly expressed in the wings (*SI Appendix*, Fig. S11).

In nymphalid butterflies, the transcription factor *optix* promotes ommochrome-based reddish coloration (Reed et al., 2011; Martin et al., 2014; Zhang et al., 2017). In *optix* knockout individuals, the expression of several ommochrome synthesis genes (e.g., *cinnabar*) is inhibited, resulting in loss of red color (Zhang et al., 2017). When we examined published RNA sequencing results from *optix* knockout individuals (Zhang et al., 2017), we found that expression of *3OHKT* was drastically reduced in *optix* knockout individuals in the pupal wings of two nymphalid butterflies, *Vanessa cardui* and *Junonia coenia* (*SI Appendix*, Fig. S12).

### *3OHKT* is essential for reddish wing pigmentation in *A. hyperbius*

Finally, to determine the contribution of 3OHKT to wing pigmentation, we performed topical RNAi using electroporation on *A. hyperbius* wings (*SI Appendix*, Fig. S13). In lepidopteran species, RNAi does not work systemically, whereas *in vivo* electroporation after small interfering RNA injection can prevent gene expression topically around the region where the positive electrode is placed (Ando & Fujiwara, 2013). RNAi of *3OHKT* caused the orange color to become pale in all individuals, while no changes were observed in the black region (Fig. 7A, B). In contrast, *cinnabar* RNAi had no effect on wing coloration in all individuals, as in *EGFP* RNAi for negative control (*SI Appendix*, Fig. S13). Meanwhile, *multicopper oxidase 2* (*MCO2*) RNAi inhibited scale development in both orange and black regions (*SI Appendix*, Fig. S13), as reported in other butterflies (Peng et al. 2020).

**Fig. 7.**
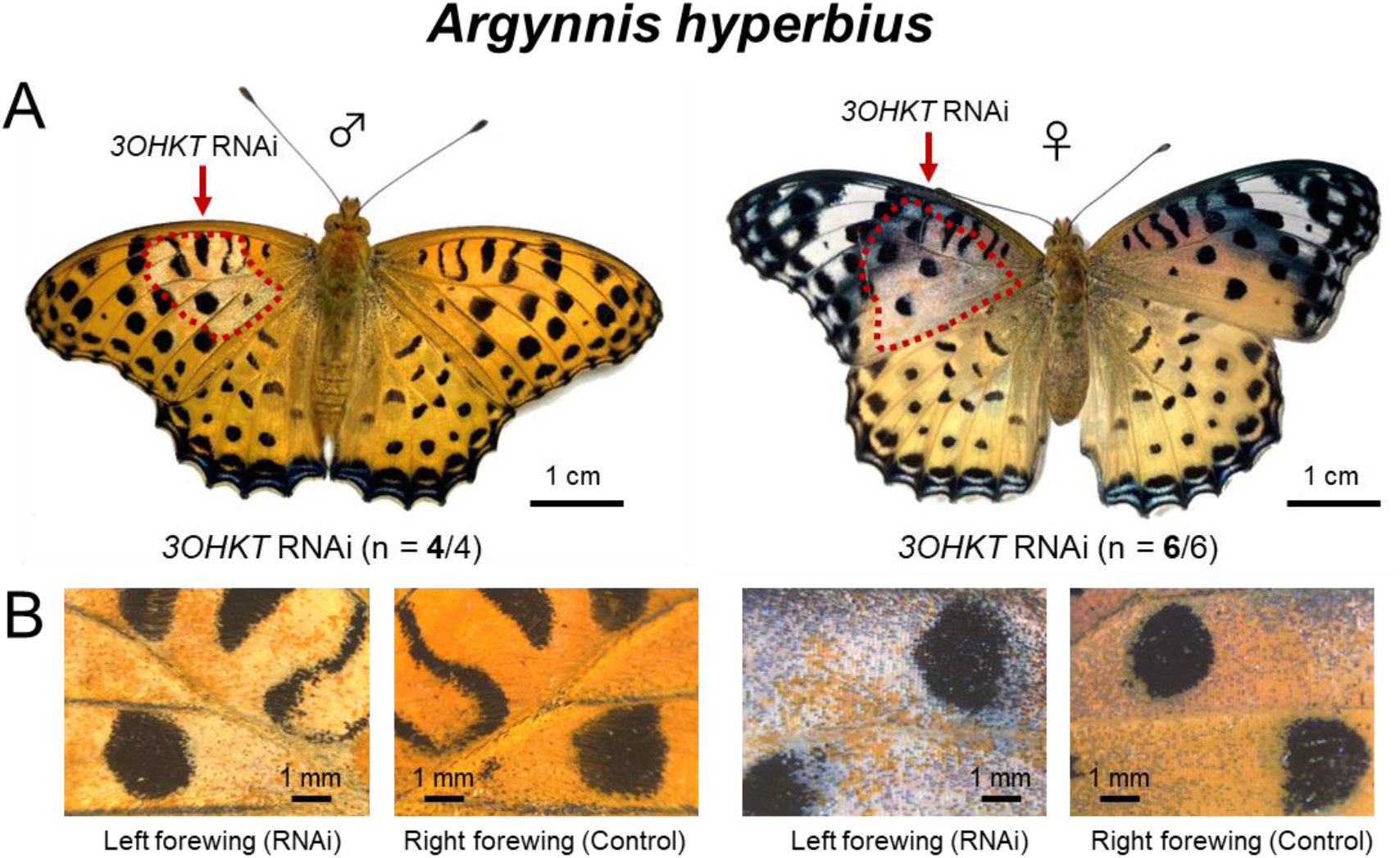
*3OHKT* RNAi phenotypes on the wings of *A. hyperbius*. (A) Whole view of male and female. Red arrows and dotted lines indicate the regions of suppressed pigmentation. Numbers in parentheses indicate (number of individuals affected by RNAi / number of emerged adults). (B) Magnified views of RNAi and control regions. See also *SI Appendix*, Figure S13.

## Discussion

### Differences in ommochrome synthesis between tissues

In this study, we elucidated that the gene responsible for the *b-t* mutant of *B. mori* is a previously-uncharacterized monocarboxylate transporter. The results of protein localization and measurement of 3OHK suggest that this gene functions as a cellular transporter for 3OHK in several tissues (Fig. 8). However, no significant differences could be detected in the color and absorbance spectra of pigment extracts of the compound eyes between wild-type and *b-t* mutant. It should be noted that results of feeding experiments of *B. mori* egg extracts to *D. melanogaster* (Kikkawa, 1941), and transplantation experiments of eye primordium of *B. mori* between wild-type and ommochrome mutants (Morohoshi, 1938) indicate that compound eye has the ability to synthesize 3OHK but also can incorporate and utilize 3OHK in the hemolymph. Considering the high expression of genes involved in the synthesis from tryptophan to 3OHK, the normal pigmentation of the compound eyes in *b-t* mutant is likely to be due to the high amount of 3OHK synthesized from tryptophan in pigment cells of the compound eyes (Fig. 8A). Since *b-t* mutant and 3OHKT deficient individuals of *B. mori* exhibited pigmentation abnormalities not only in eggs but also in larval epidermis and ganglions, and *3OHKT* RNAi prevented normal orange pigmentation in wings of *A. hyperbius*, 3OHK uptake is necessary for ommochrome-based pigmentation in tissues other than compound eyes (Fig. 8B, 8C).

**Fig. 8.**
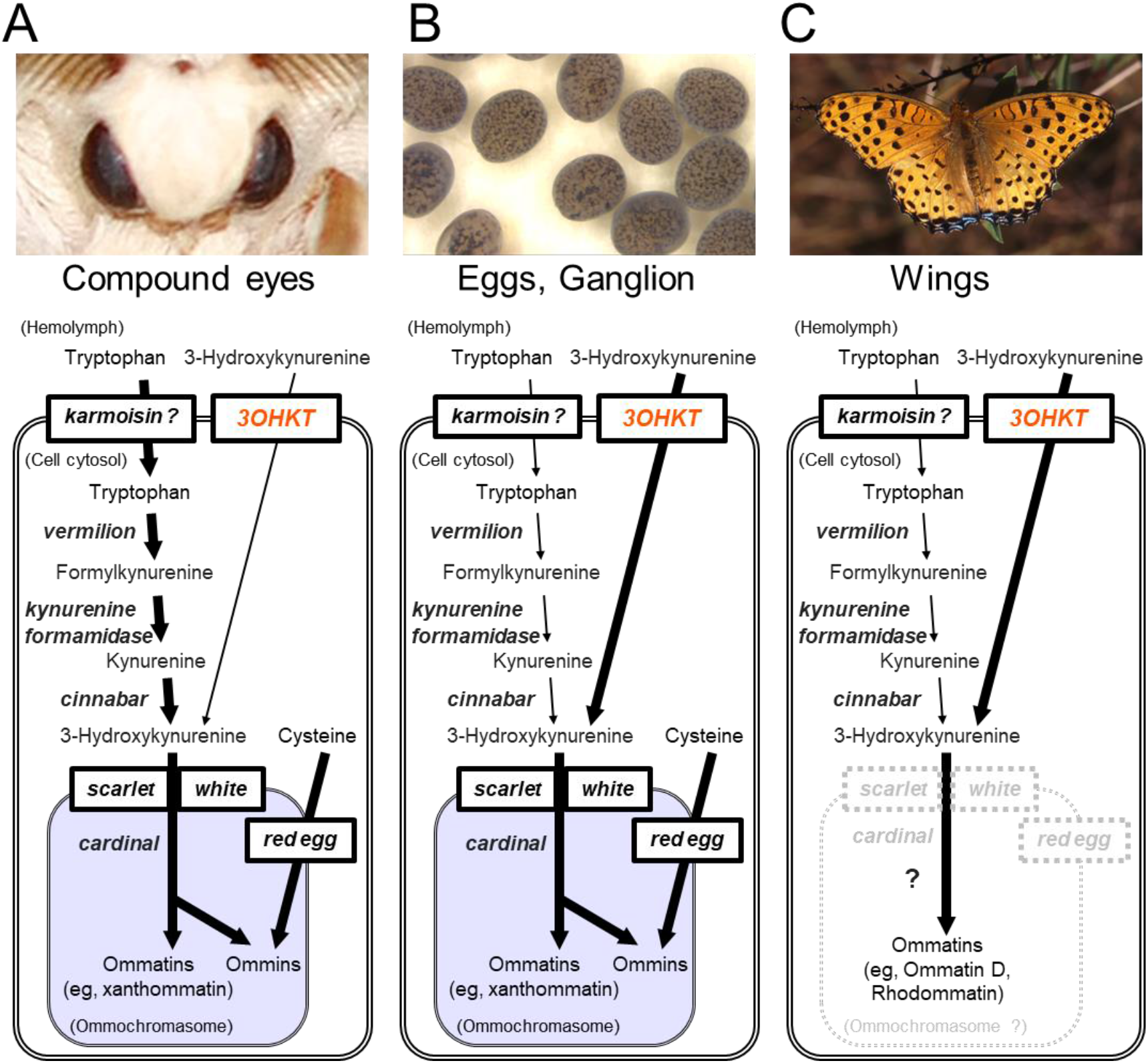
A proposed model of tissue-dependent ommochrome synthesis pathways. Examples of the compound eyes of the silkworm (A), eggs and ganglion of the silkworm (B), and the wings of nymphalid butterflies (C) are shown. Genes involved in ommochrome synthesis pathway is indicated in italic. The compound eyes of *Drosophila* are probably identical to the compound eyes of the silkworm except for the absence of *re* gene. The major differences are the contribution of 3OHKT (A vs B & C) and the presence or absence of the omochromasome (A & B vs C).

### Possible source of 3OHK in hemolymph

In the fruit fly *D. melanogaster*, the cellular uptake of tryptophan from the hemolymph has been assumed to be mediated by the major facilitator superfamily transporter gene *karmoisin* (Sullivan et al., 1974; Tearle, 1991). In *B. mori*, genes involved in the early step of ommochrome synthesis from tryptophan to 3OHK (i.e., *karmoisin, vermilion, kynurenine formamidase*, and *kynurenine monooxygenase* (= *cinnabar*) were expressed in fat body, which is not pigmented by ommochrome, as well as early embryo and brain which accumulate ommochrome pigments, according to RNAseq data deposited in silkbase (*SI Appendix*, Fig. S14, http://silkbase.ab.a.u-tokyo.ac.jp/cgi-bin/news.cgi). On the other hand, expression of genes involved in the late step of ommochrome synthesis in ommochromasomes, such as *red egg, white, cardinal* was detected in early embryo and brain, but little or no expression was detected in the fat body. Thus, 3OHK synthesized in the fat body is likely to be a major source of 3OHK in the hemolymph.

### 3OHKT-dependent ommochrome pathway should contribute to pigmentation in butterfly wings

One of the major findings in this study was that *3OHKT* was involved in ommochrome-based pigmentation of butterfly wings (Fig. 7). No effect was detected by *cinnabar* RNAi (*SI Appendix*, Fig. S13), at least in *A. hyperbius* wings, suggesting that uptake from hemolymph rather than *de novo* synthesis is the main source of 3OHK in wings (Fig. 8C). This result is consistent with previous findings that 3OHK is more efficiently incorporated into pupal wings than tryptophan in *A. levana* (Koch, 1993). Notably, genes involved in the late step of ommochrome synthesis in the granules called ommochromasomes (e.g., *cardinal, red egg, scarlet*) were scarcely expressed in the wings (*SI Appendix*, Figs. S10, S11), which is consistent with classical reports that ommatin D and rhodommatin do not form granules (Linzen, 1974). These results suggest that the ommochrome synthetic pathway differs greatly between wings (Fig.8C) and the compound eyes (Fig.8A).

Previous studies of ommochrome-related genes have focused primarily on the synthesis of 3OHK from tryptophan such as *vermilion* and *cinnabar*, or on the transport of 3OHK into ommochromasome such as *scarlet* and *white* (Reed and Nagy, 2005; Reed et al., 2008; Ferguson and Jiggins, 2009; Matsuoka and Monteiro, 2018). Notably, in the nymphalid butterfly *Bicyclus anynana*, the gene disruption of *vermilion* by CRISPR/Cas9 did not show any phenotypes in G0 generation, which is consistent with our proposed 3OHKT-dependent ommochrome pathway in wings (Fig.8C). Moreover, gene disruption of *white* or *scarlet* showed reduced pigmentation in compound eyes and/or larval epidermis, but not in adult wings (Matsuoka and Monteiro, 2018), suggesting that omochromasome-independent pigmentation occurs in the wings (Fig.8C). To elucidate the ommochrome synthesis pathway in butterfly wings, the candidate genes involved in the downstream of 3OHK should be functionally investigated in the future.

### Materials and methods

The *b-t* mutant (e30 strain) of *B. mori* was provided from the silkworm stock center of Kyushu University supported by the National BioResource Project. The wild-type silkworm strain p50T was maintained in the Silkworm Research Group, The National Agriculture and Food Research Organization (NARO), and other silkworm strains C108, *pnd* were maintained in The Research Center of Genetic Resources, NARO. Silkworms were reared on an artificial diet (Nihon Nosan Kogyo) under a 12-h-light: 12-h-dark photoperiod at 28°C from first to fourth instar, and at 25°C from fifth instar. Final instar larvae of *A. io* were collected at Ikeda-cho, Hokkaido, and larvae of *A. hyperbius* were collected at Tsukuba, Ibaraki, or Setagaya, Tokyo.

In order to conduct ddRAD-seq and mapping of the *b-t* locus, F_1_ individuals were obtained from a single cross between the *b-t* mutant (e30 strain) female (*b-t* / *b-t*) and the wild-type (C108) male (+ ^*b-t*^ / + ^*b-t*^). F_1_ male moths (*b-t* / + ^*b-t*^) were then crossed with *b-t* female moths (*b-t* / *b-t*) to obtain BC1 eggs. The genotypes of BC1 females (*b-t* / *b-t* or *b-t* / + ^*b-t*^) were determined by the color of eggs they laid. Genomic DNAs were extracted from 38 *b-t* (*b-t* / *b-t*) BC1 females, 52 + ^*b-t*^ BC1 females (*b-t* / + ^*b-t*^), and their parents, using the Maxwell 16 Tissue DNA Purification kit (Promega). The ddRAD-seq library construction, sequencing, and single nucleotide polymorphisms (SNPs) extraction were conducted according to Tomihara et al., 2021.

For RNAi experiments, small interfering RNA (siRNA) was designed using the siDirect program (v.2.0; http://sidirect2.rnai.jp/) (Naito et al., 2009). The sequences of the siRNA are listed in Table S1. (*SI Appendix*). siRNA for the *enhanced green fluorescent protein* (*EGFP*) gene used as a control is the same with Ando and Fujiwara 2013. 1 μL of 100 μM siRNA solution was injected into the base of the left forewing of day 0 pupae using a glass needle. Immediately after injection, platinum electrodes and droplets of conductive gel (Ultrasound GELH-250, Nepagene, Ichikawa, Japan) were placed nearby and fifteen pulses of 25 V (each 280 ms pulse/s) were applied by the electroporator Cure-Gene (CellProduce Co., Ltd, Tokyo, Japan) (*SI Appendix*, Fig. S13A).

Complete details on the materials and methods are available in SI Appendix.

## Supporting Information

Materials/Methods, Supplementary Text, Tables, Figures, and References

## Data Availability

DNA sequences and raw Fastq data have been deposited in DNA Data Bank Japan Read Archive (DRA011673). All other study data are included in the article and/or supporting information.

## Acknowledgements

We thank Yuusuke Kobayashi for helpful advice on phylogenetic analyses, Takumi Suzuki for allowing us to use his equipment, Ryuunosuke Chiba for initial investigation of mutant phenotype analysis, Yasuhito Yagihashi and Haruka Suzuki for technical support for immunocytochemistry, Jun Okude for collecting *Aglais io* larvae, and Bin Hirota, Minoru Osanai, Naoki Futahashi for collecting *Argyreus hyperbius* larvae. We are grateful to Kaoru Nakamura, Toshihiko Misawa, Haruna Hirota for rearing silkworms and Kaoru Nakamura, Toshihiko Misawa for technical support on silkworm embryonic injection. This work was supported by JSPS KAKENHI Grant Number 26850220 and Ibaraki University Grant for Specially Promoted Research to M. O-F, Cooperative Research Grant of the Genome Research for BioResource, NODAI Genome Research Center, Tokyo University of Agriculture, and the National BioResource Project, Japan.

## Author contributions

M.O.-F. designed research, Hirosumi U., M.O.-F., M.M., Y.T., R.F., G.O., K.U., T.I., Hironobu U., S.Y., Y.B., and T.T. performed research; H.S., and K. Yamamoto contributed new reagents/analytic tools; K. Yoshitake, M.O.-F., R.F., Hirosumi U analyzed data; MO-F, Hirosumi U, R.F., M.M., G.O. wrote the paper. All authors approved the final version of the manuscript and agree to be accountable for all aspects of the work.

## References

T. Ando, H. Fujiwara, Electroporation-mediated somatic transgenesis for rapid functional analysis in insects. Development 140, 454–458 (2013).

C. S. Brent, J. J. Hull, RNA interference-mediated knockdown of eye coloration genes in the western tarnished plant bug (Lygus hesperus Knight). Arch. Insect Biochem. Physiol. 100, e21527 (2019).

G. Broehan, T. Kroeger, M. Lorenzen, H. Merzendorfer, Functional analysis of the ATP-binding cassette (ABC) transporter gene family of Tribolium castaneum. BMC Genomics 14, 6 (2013).

H. Doira, H. Kihara, H. Chikushi, R. Umezu, Genetical studies on the “maternal brown of Tsujita” mutation in Bombyx mori. J. Sericultural Sci. Japan 50, 154–157 (1981). In Japanese.

L. C. Ferguson, C. D. Jiggins, Shared and divergent expression domains on mimetic Heliconius wings. Evol Dev. 11, 498–512 (2009).

F. Figon, J. Casas, Ommochromes in invertebrates: biochemistry and cell biology. Biol. Rev. 94, 156–183 (2019).

R. Futahashi, M. Osanai-Futahashi, “Pigments in Insects”, in Pigments, Pigment Cells and Pigment Patterns, H. Hashimoto, M. Goda, R. Futahashi, R. Kelsh, T. Akiyama, Eds. (Springer, 2021) pp. 3–43.

N. Grubbs, S. Haas, R. W. Beeman, M. D. Lorenzen, The ABCs of eye color in Tribolium castaneum: Orthologs of the Drosophila white, scarlet, and brown genes. Genetics 199, 749–759 (2015).

D. A. Harris, K. Kim, K. Nakahara, C. Vásquez-Doorman, R. W. Carthew, Cargo sorting to lysosome-related organelles regulates siRNA-mediated gene silencing. J. Cell Biol. 194, 77–87 (2011).

H. Kikkawa, Mechanism of Pigment Formation in Bombyx and Drosophila. Genetics 26, 587–607 (1941).

P. B. Koch, Precursors of pattern specific ommatin in red wing scales of the polyphenic butterfly Araschnia levana L.: haemolymph tryptophan and 3-hydroxykynurenine. Insect Biochem 21, 785–794 (1991)

P. B. Koch, Production of [14C]-labeled 3-hydroxy-L-kynurenine in a butterfly, Heliconius charitonia L. (Heliconidae), and precursor studies in butterfly wing ommatins. Pigment Cell Res 6, 85–90 (1993)

N. Kômoto, G.-X. Quan, H. Sezutsu, T. Tamura, A single-base deletion in an ABC transporter gene causes white eyes, white eggs, and translucent larval skin in the silkworm w-3^oe^ mutant. Insect Biochem. Mol. Biol. 39, 152–156 (2009).

B. Linzen, The Tryptophan → Omrnochrome Pathway in Insects. Adv. In Insect Phys. 10, 117–246 (1974).

M. D. Lorenzen, S. J. Brown, R. E. Denell, R. W. Beeman, Cloning and characterization of the Tribolium castaneum eye-color genes encoding tryptophan oxygenase and kynurenine 3-monooxygenase. Genetics 160, 225–234 (2002).

A. Martin, K.J. McCulloch, N.H. Patel, A.D. Briscoe, L.E. Gilbert, R.D. Reed, Multiple recent co-options of Optix associated with novel traits in adaptive butterfly wing radiations. Evodevo, 5, 7 (2014).

Y. Matsuoka, A. Monteiro, Melanin pathway genes regulate color and morphology of butterfly wing scales. Cell Rep. 24, 56–65 (2018).

S. Morohoshi, Transplantation of optic discs and eye colours in Bombyx mori. Japanese J. Genet. 14, 204–210 (1938) In Japanese.

Y. Naito, J. Yoshimura, S. Morishita, K. Ui-Tei, siDirect 2.0: updated software for designing functional siRNA with reduced seed-dependent off-target effect. BMC Bioinformatics 10, 392 (2009).

H. F. Nijhout, The Development and Evolution of Butterfly Wing Patterns, (Smithsonian Institution Scholarly Press, 1991).

M. Osanai-Futahashi et al., Identification of the Bombyx red egg gene reveals involvement of a novel transporter family gene in late steps of the insect ommochrome biosynthesis pathway. J. Biol. Chem. 287, 17706–17714 (2012).

M. Osanai-Futahashi et al., Positional cloning of a Bombyx pink eyed white egg locus reveals the major role of cardinal in ommochrome synthesis. Heredity, 116, 135–145 (2016).

C. L. Peng, A. Mazo-Vargas, B. J. Brack, R. D. Reed, Multiple roles for laccase2 in butterfly wing pigmentation, scale development, and cuticle tanning. Evol Dev. 22: 336–341 (2020).

M. Pepling, S. M. Mount, Sequence of a cDNA from the Drosophila melanogaster white gene. Nucleic Acids Res. 18, 1633 (1990).

G. X. Quan et al., Rescue of mutant by introduction of the wild-type Bombyx kynurenine 3-monooxygenase gene. Insect Sci. 14, 85–92 (2007).

R. D. Reed, W. O. McMillan, L. M. Nagy, Gene expression underlying adaptive variation in Heliconius wing patterns: non-modular regulation of overlapping cinnabar and vermilion prepatterns. Proc Biol Sci. 275: 37–45 (2008).

R. D. Reed, L. M. Nagy, Evolutionary redeployment of a biosynthetic module: Expression of eye pigment genes vermilion, cinnabar, and white in butterfly wing development. Evol. Dev. 7, 301–311 (2005).

R. D. Reed, R. Papa, A. Martin, H M. Hines, B. A. Counterman, C. Pardo-Diaz, C. D. Jiggins, N. L. Chamberlain, K. R. Kronforst, R. Chen, G. Halder, H. F. Nijhout, W. O. McMillan. optix drives the repeated convergent evolution of butterfly wing pattern mimicry. Science, 333, 1137–1141 (2011).

H. Sonobe, E. Ohnishi, Accumulation of 3-hydroxykynurenine in ovarian follicles in relation to diapause in the silkworm, Bombyx mori L. Dev. Growth Differ. 12, 41–52. (1970).

L. L. Searles, R. S. Ruth, A. M. Pret, R. A. Fridell, A. J. Ali, Structure and transcription of the Drosophila melanogaster vermilion gene and several mutant alleles. Mol. Cell. Biol. 10, 1423–1431 (1990).

L. L. Searles, R. A. Voelker, Molecular characterization of the Drosophila vermilion locus and its suppressible alleles. Proc. Natl. Acad. Sci. 83, 404–408 (1986).

D. T. Sullivan, S. L. Grillo, R. J. Kitos, Subcellular localization of the first three enzymes of the ommochrome synthetic pathway in Drosophila melanogaster. J. Exp. Zool. 188, 225–233 (1974).

K. Tatematsu et al., Positional cloning of silkworm white egg 2 (w-2) locus shows functional conservation and diversification of ABC transporters for pigmentation in insects. Genes to Cells 16, 331–342 (2011).

R. Tearle, Tissue specific effects of ommochrome pathway mutations in Drosophila melanogaster. Genet. Res. B, 257–266 (1991).

R. G. Tearle, J. M. Belote, M. McKeown, B. S. Baker, A. J. Howells, Cloning and characterization of the scarlet gene of Drosophila melanogaster. Genetics 122, 595–606 (1989).

K. Tomihara et al., 2021. Mutations in a β-group of solute carrier gene are responsible for egg and eye coloration of the brown egg 4 (b-4) mutant in the silkworm, Bombyx mori. Insect Biochem. Mol. Biol. 137, 103624.

L. Wang et al., Bombyx mori monocarboxylate transporter 9 (BmMCT9) is involved in the transport of uric acid in silkworm integument. Genes to Cells 25, 33–40 (2020).

W. D. Warren, S. Palmer, A. J. Howells, Molecular characterization of the cinnabar region of Drosophila melanogaster: Identification of the cinnabar transcription unit. Genetica 98, 249–262 (1996).

T. L. Williams et al., Dynamic pigmentary and structural coloration within cephalopod chromatophore organs. Nat. Commun. 10, 1004 (2019).

L. Zhang, A. Mazo-Vargas, R.D. Reed, Single master regulatory gene coordinates the evolution and development of butterfly color and iridescence. Proc Natl Acad Sci U S A. 114, 10707–10712 (2017).

