## Supplementary material for "A novel monocarboxylate transporter involved in 3-hydroxykynurenine transport for ommochrome coloration": SI Appendix

### SI Materials and Methods

#### Cell Culture

The silkworm cell line BmN4-SID1 (Mon et al., 2012), a gift from Dr. Kusakabe, Kyushu University. The cells were subcultured once a week by exchanging medium with 1/5 dilution, and BmN4-SID1 cells were maintained at 27℃in IPL41 medium (Gibco) supplemented with 10% fetal bovine serum (Biosera).

#### RNA sequencing (RNA-seq)

Total RNA was isolated from eggs of the silkworm *Bombyx mori* at 0, 24, 48, and 72 h post oviposition, or from pupal wings and eyes of the nymphalid butterflies *Aglais io* and *Argynnis hyperbius* using a Maxwell 16 LEV Simply RNA Tissue kit (Promega). RNA sequencing was performed as described previously (Okude et al., 2022). Using 1 µg of total RNA per sample as template, cDNA libraries were constructed using TruSeq RNA Sample Preparation Kits v2 (Illumina), TruSeq Stranded mRNA Library Prep Kit (Illumina), or NEBNext Ultra II Directional RNA Library Prep Kit for Illumina (New England Biolabs) and sequenced by HiSeq (Illumina). We mapped the RNA-seq reads of *B. mori* to the predicted genes set of *B. mori* (Kawamoto et al., 2019) with the Salmon program (version 1.5.1) (Patro et al., 2017). For *A. io* and *A. hyperbius,* RNA-seq reads were subjected to *de novo* assembly using the Trinity program v. 2.4.0 (Grabherr et al., 2011). After automatic assembling, we checked and manually corrected the sequences of genes involved in ommochrome synthesis pathway using the Integrative Genomics Viewer (Robinson et al., 2011). After revising the sequence, mapping was performed using the Salmon program (version 1.5.1) (Patro et al., 2017). Raw fastq data have been deposited in DDBJ under accession number DRA011673. The sample information for RNA sequencing is shown in *SI Appendix*, Table S2.

#### ddRAD-seq

The ddRAD-seq library was constructed according to the method described in Tomihara et al., (2021). EcoRI and MspI was used for double digestion of genomic DNAs. Sequencing was conducted using an Illumina HiSeq 2500, generating 200-bp paired-end reads.

#### Wave scanning of pigments by spectrophotometer

Adult compound eye and egg pigment extracts were prepared from strains C108, e30, and 3OHKT Δ10, respectively. For each strain, ten eggs each from four batches were homogenized by plastic pestle in 150 µL of 5% hydrochloric acid in methanol (v/v) to extract ommochrome pigments. For compound eye pigment extraction, an adult compound eye each from four individuals were used for each strain. The extracts were vortexed for 10 min and centrifuged at 25℃ 11000 × g for 1 min, and the supernatants were collected and used for analyses. Oxidizing agent (1% NaNO2) and reducing agent (1% ascorbic acid) were added to every extract to observe redox-dependent color changes. Wave scanning of samples was performed using a GeneQuant 1300 spectrophotometer (Biochrom) in the range of 300 nm to 600 nm.

#### Transmembrane prediction

Transmembrane prediction was performed using Phobius (Käll et al., 2007).

#### Genomic PCR for detecting genomic deletion of b-t

Genomic DNA was exracted from adult legs using DNAzol (Molecular Research Center, Inc.), and PCR was conducted with KOD one (TOYOBO) using primers b-t_intron-F7 and b-t_exon9-R2 (*SI Appendix*, Table S3).

#### *TALEN* mediated gene knockout

TALEN knockout experiments were conducted as reported previously (Takasu *et al*, 2013). A manual search was performed for a TALEN target site for exon 9 of *3OHKT*. TCAGGGGTCCTCTACG and TGCCAAGAAACAATACACGT in *3OHKT*, were selected as the recognition sequences. The TAL portions were prepared by Golden Gate TALEN and TAL Effector Kit (Addgene), following the method described by Cermak et al, (2011), using an in-vitro expression vector pBlue-TAL. The constructed plasmids were purified by HiSpeed Plasmid Midi kit (Qiagen), linearized by XbaI, treated with proteinase K, extracted with phenol-chloroform-isoamylalcohol (25:24:1) and chloroform, precipitated with ethanol, and washed with 70% ethanol three times. TALEN mRNAs were transcribed by mMESSAGE mMACHINE kit (Applied Biosystems), followed by lithium chloride precipitation, and were then washed with 70% ethanol three times. The resulting TALEN mRNAs were dissolved into injection buffer (0.5 mM phosphate buffer [pH 7.0], 5 mM KCl) to achieve a final concentration of 0.5 mg/mL each, and were microinjected into 192 eggs of non-diapausing *B. mori pnd* (wild-type egg color) strain 3–5 h after laying.

Of the 192 injected eggs, 158 became adults. 78 G0 males were mated with sibling G0 females to obtain G1 generation eggs. For founder brood screening, one fourth of the larvae from each G1 brood were pooled, and genomic DNA was extracted using DNAzol (Molecular Research Center, Inc.). The TALEN targeted region was amplified by PCR with primers Bmb-t_wt-genotyping-F and Bmb-t_wt-genotyping-R (*SI Appendix*, Table S3), and PCR products were subjected to heteroduplex assay. The remaining larvae of the brood that was judged to have high mutation rate from the heteroduplex assay was reared sib-mated. For genotyping G1 and G2 individuals, genomic DNA was extracted from adult legs using DNAzol, and the targeted region was amplified by PCR with primers Bmb-t_wt-genotyping-F and Bmb-t_wt-genotyping-R (*SI Appendix*, Table S3), and sequenced by ABI prism 3130 Genetic Analyzer (Applied Biosystems).

To confirm that *3OHKT* gene corresponds to the *b-t* locus, complementation test was performed by crossing *b-t* (e30 strain) mutant females with and *3OHKT* knockout G2 males Δ5+I2/Δ4 and Δ10. The next generation was raised and dissected for inspection of ganglia pigmentation at the larval stage or sib-mated to check the color of the eggs.

To obtain homozygous strain of the knockout allele, the G2 generation was outcrossed with wild-type (C108), and their F1 offsprings were sib-mated. Genotype of F1 individuals were checked by PCR and sequencing the genome extracted from adult legs. To select *3OHKT* knockout homozygotes from the F2 generation, first instar larvae with pale colored abdomen were selected from F2 broods which both parents have the same genotype, and intercrossed. As a result, *3OHKT* Δ10 homozygous strain was established, which laid the similar pale brown color eggs as *b-t* (e30 strain).

**Phylogenetic analysis**

We performed a phylogenetic analysis using protein sequences from 13 insects (*B. mori, Helicoverpa armigera, Papilio polytes, Drosophila melanogaster, Aedes aegypti, Anopheles gambiae, Apis mellifera, Bombus impatiens, Harpegnathos saltator, Tribolium castaneum, Onthophagus taurus, Cimex lectularius, Blattella germanica, Zootermopsis nevadensis*), and 6 crustaceans (*Armadillidium vulgare, Chionoecetes opilio, Darwinula stevensoni, Penaeus monodon, Penaeus vannamei, Portunus trituberculatus*), that showed hits to *B. mori* 3OHKT with significance of E-value < $e^{-19}$, and monocarboxylate transporter proteins retrieved from homology search to RNAseq data of pupal wings and compound eyes of peacock butterfly *Aglais io*. Deduced amino acid sequences of *3OHKT* gene homologs were aligned using the MUSCLE program (Edgar, 2004) installed in MEGA6 (Tamura *et al*, 2013). Poorly aligned regions was removed by TrimAl (Capella-Gutiérrez *et al*., 2009). Molecular phylogenetic analysis was conducted by neighbor-joining methods using MEGA6 and Maximum-likelihood methods using RAxML (Stamatakis, 2014). The evolutionary distances for Neighbor-joining and Maximum-likelihood were computed using the JTT model, and a LG+F Gamma model that was selected by Aminosan (Tanabe, 2011), respectively. The confidence of the phylogenetic lineages was assessed by bootstrap analysis.

**Plasmid construction and transfection**

The full-length coding regions of *3OHKT* was amplified by PCR using P50T ovary cDNA as template. The primers used are listed in *SI Appendix*, Table S3. The PCR products were inserted into the pBac-pIZ-N-3xFlag or pBac-pIZ-C-3xFlag (Suzuki et al., in preparation) by HiFi Assembly method (New England Biolabs). Plasmids were transfected into BmN4-SID1 cells using TransIT insect transfection Regent (Mirus). The cells were cultured 3 days at 25°C before immunostaining.

**Immunocytochemistry**

The cells transfected with plasmids were transferred to coverslips coated with 30 μL of 0.35 mg/ml Concanavalin A 1 hour before fixation. The coverslip was washed 3 times with PBS, and fixed with 1 mM EGTA in MeOH at -30℃ for 2 min. The fixed cells were washed 3 times with PBST and blocked with 10% goat serum in PBST at room temperature for 1 hour. After blocking, a primary antibody reaction was carried out using mouse monoclonal-anti FLAG (SIGMA Product Number F1804) at a dilution of 1:200 for overnight at 4℃. After three washes, secondary antibody reaction was carried out with goat anti-Mouse IgG (H+L) highly Cross-adsorbed Secondary Antibody, Alexa Fluor 555 (Invitrogen, A21412) at a dilution of 1:1000 for 1 hour at room temperature. After three washes, the cells were mounted with Vectashield mounting medium with DAPI (VECTOR). Images were acquired on OLYMPUS IX81 FV1000 Laser Scanning Confocal Microscope.

**Quantification of 3-hydroxykynurenine**

Strains C108 (WT), e30 (*b-t*), *3OHKT*Δ10 were used for 3-hydroxykynurenine quantification. Eggs were collected at 24h post oviposition, and hemolymph was collected from day 4 female pupae. Ten eggs homogenized in 1000 μL of methanol containing 0.1% (v/v) formic acid (acidified methanol), while 20 μL of hemolymph was resuspended to 180 μL the same solvent. They were subjected to ultrasonication for 3 min and centrifugation for 10 min, and the supernatant was collected. The remnants were re-extracted with 200 μL of acidified methanol. 100 μL aliquot of the combined extract was dried in vacuo, resuspended to 100 μL of water containing 0.1% (v/v) formic acid, and loaded on a reversed-phase solid-phase extraction column (GL-Tip SDB, GL Science) for further purification. The flow-through fraction was collected and analyzed using a liquid chromatography-mass spectrometry (LC-MS) system. This system consisted of an ultrahigh-performance liquid chromatograph equipped with a photodiode array detector (Acquity UPLC H-Class, Waters) and a quadrupole-time-of-flight mass spectrometer with an electrospray ionization source (Xevo-G2XS, Waters). The 3-hydroxykynurenine was separated on an ACQUITY UPLC BEH C18 column (2 mm i.d. x 100 mm length, Waters) under a water-acetonitrile gradient regime at a flow rate of 0.25 mL/min. While a chromatographic peak responsible for 3-hydroxykynurenine was identified based on the absorbance spectrum and MS/MS fragment spectrum of a standard reagent (Merck), absorbance peak area at 372 nm was used for quantification.

**RNA isolation and quantitative RT-PCR**

Total RNA were isolated from ovaries and compound eyes of wild-type strain (C108) at pupa day 4 and 8, using ISOGENⅡ(Nippon Gene) according to the manufacturer's instructions, and reverse transcribed with a random primer ($N_6$) using the First-Stand cDNA Synthesis Kit (Cytiva).

To estimate the abundance of *3OHKT* transcripts, real-time PCR was carried out using Step One™ (Applied Biosystems) Software v 2.3 by the relative standard curve method. PCR reaction was conducted using TB Green Rremix ExTaq Ⅱ (TaKaRa) with the primer sets listed in *SI Appendix*, Table S3. The *ribosomal protein L3* (*rPL3*, GenBank:AY769270.1) was used as an internal control.

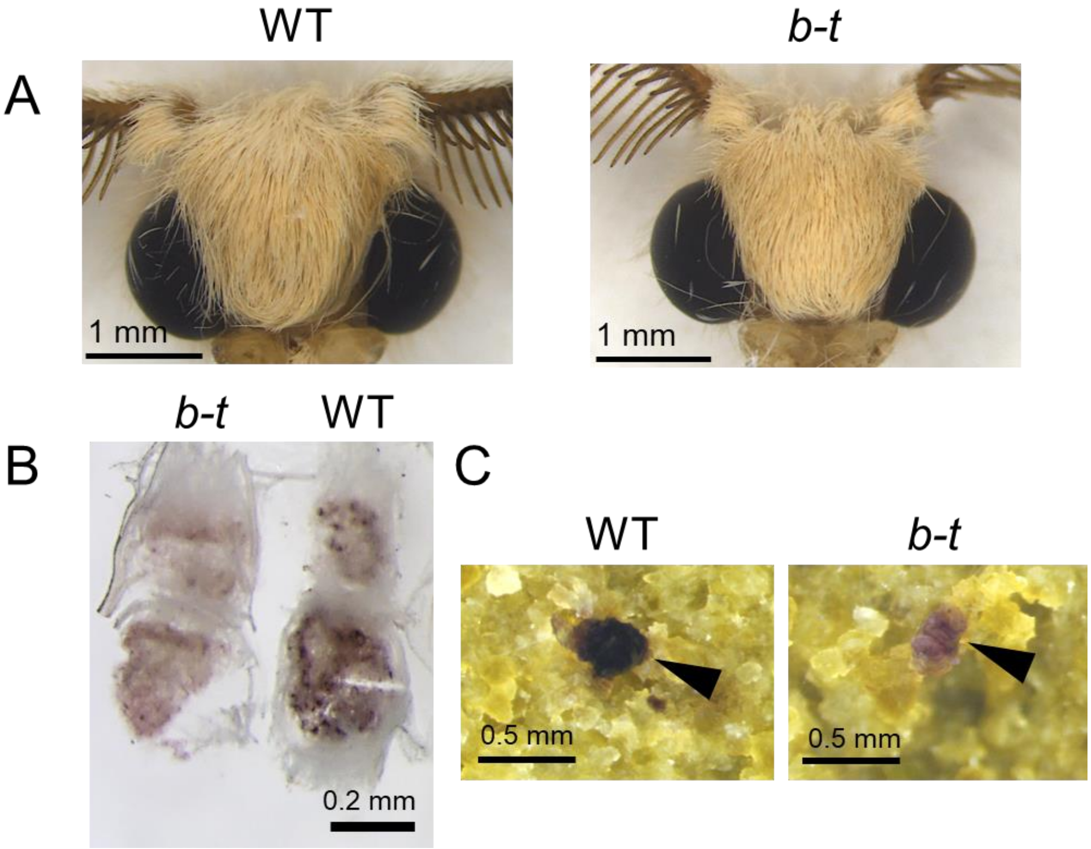


**Fig. S1. Wild-type and *b-t* mutant of the silkworm *B. mori.***
(A) Adult compound eyes of wild-type (C108 strain) and *b-t* (e30 strain). (B) The fifth-larval ganglion of *b-t* (e30 strain)and wild-type (C108 strain). (C) Feces of wild-type (C108 strain) and *b-t* (e30 strain).

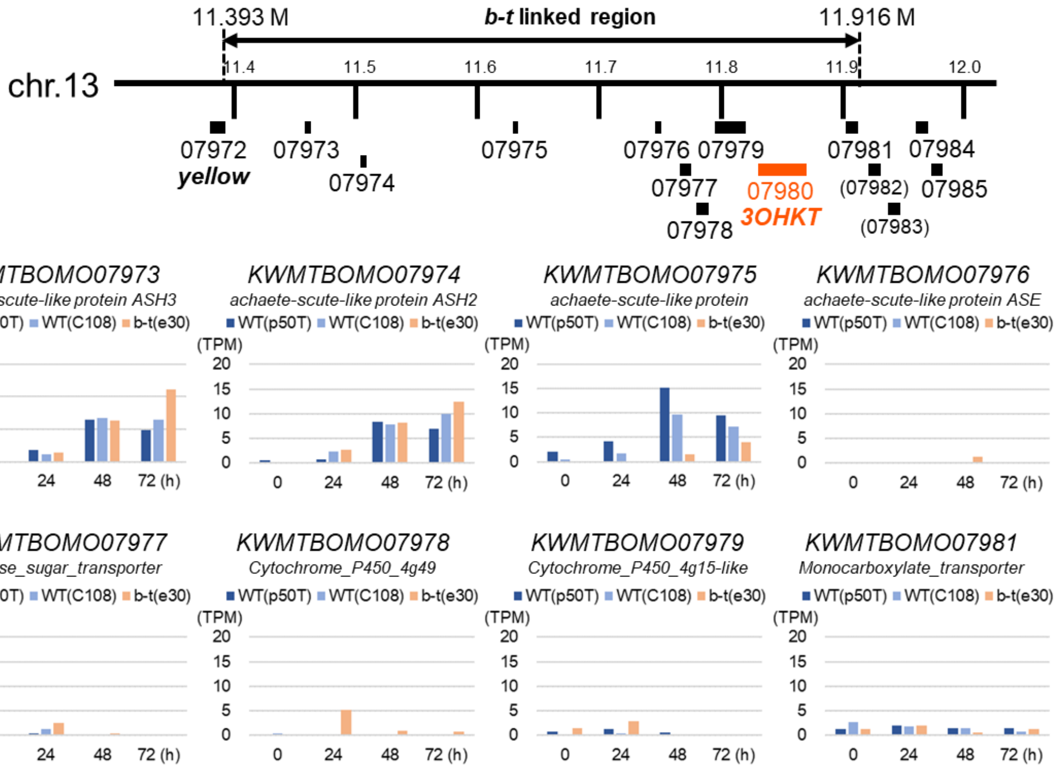


**Fig. S2. Expression analysis of genes within *b-t*-linked region of *B. mori*.**
The expression data of *KWMTBOMO07980* (*3OHKT*) is shown in Fig. 2C.

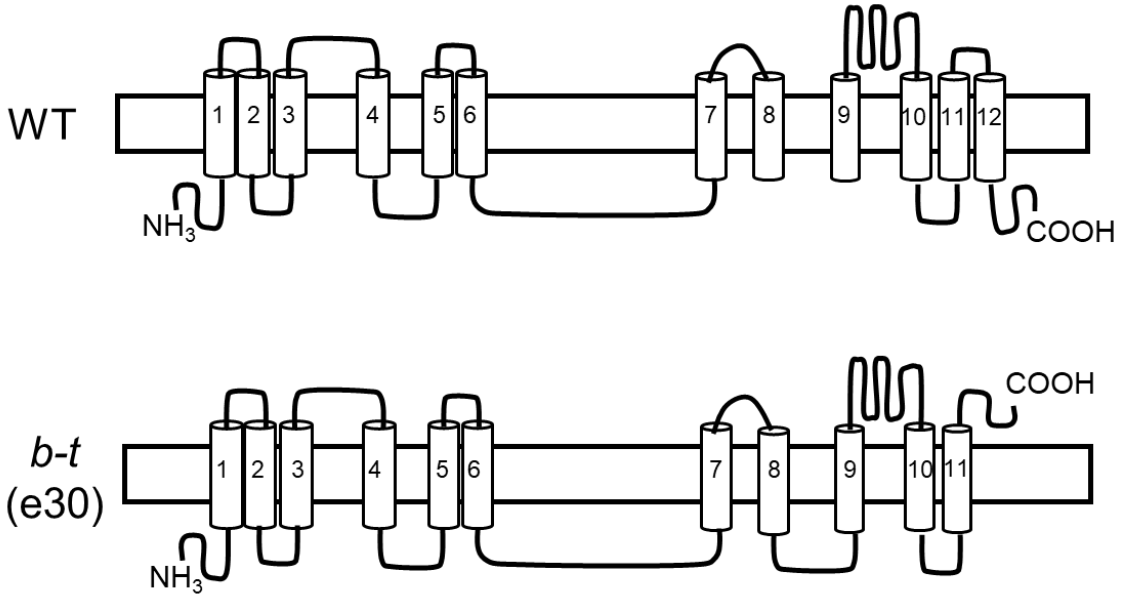


**Fig. S3. Transmembrane domain predictions of *KWMTBOMO07980* (*3OHKT)* gene by Phobius.**
Numbers indicate the transmembrane helices.

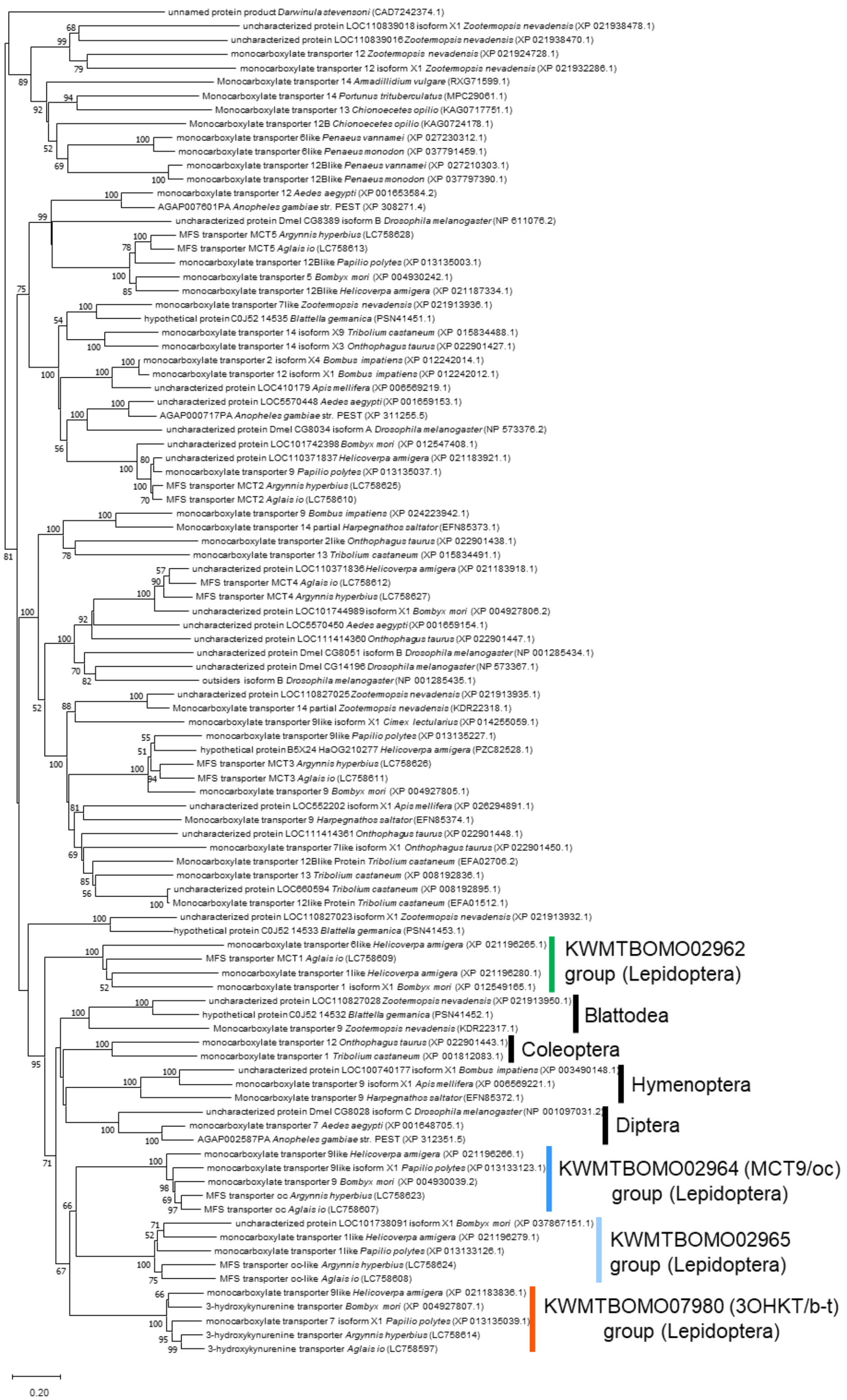


**Fig. S4. Phylogenetic relationship of *KWMTBOMO07980* (*3OHKT*) gene homolog in insects.** Phylogenetic tree constructed based on neighbor-joining method. Bootstrap values are indicated on each node. Pale orange vertical line indicates lepidopteran 3OHKT orthologs. Pale blue, navy and green vertical lines indicates lepidopteran paralogs of *B. mori* 3OHKT.

A

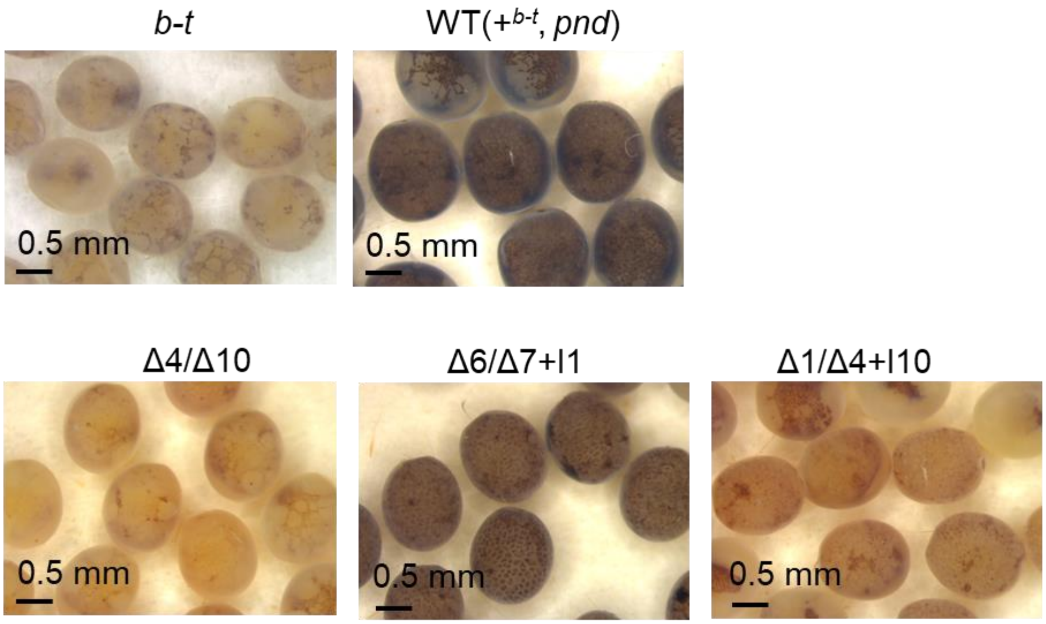


B

| | sequence | Indel |
|---|---|---|
| Wild-type | TTTTTTCAGGGGTCCTCTACGAAGAAACAGGCAGCTACGTGTATTGTTTCTTGGCAATGAGTTTTT | none |
| G1 female #4-1 | TTTTTTCAGGGGTCCTCTACGAAGAAACAG----CTACGTGTATTGTTTCTTGGCAATGAGTTTTT | Δ4 |
|  | TTTTTTCAGGGGTCCTCTACGAAGAAAC----------GTGTATTGTTTCTTGGCAATGAGTTTTT | Δ10 |
| G1 female #4-2 | TTTTTTCAGGGGTCCTCTACGAAGAAA------GCTACGTGTATTGTTTCTTGGCAATGAGTTTTT | Δ6 |
|  | TTTTTTCAGGGGTCCTCTACGAAGAAG------GCTACGTGTATTGTTTCTTGGCAATGAGTTTTT | Δ7+I1 |
| G1 female #19-2 | TTTTTTCAGG-GTCCTCTACGAAGAAACAGGCAGCTACGTGTATTGTTTCTTGGCAATGAGTTTTT | Δ1 |
| | TTTTTTCAGGGGTCCTCTACGAAGAAACGTGTATTGTTAGCTACGTGTATTGTTTCTTGGCAATGAGTTTTT | Δ4+I10 |

**Fig. S5. Disruption of the *KWMTBOMO07980* (*3OHKT*) gene using designed TALENs in *B. mori*.**
(A) Reprensentative photos of G2 eggs laid by *3OHKT* knockout G1 moths. The genotype of *3OHKT* of the mother moth is indicated above the photos. *b-t* (e30 strain), and wild-type (+$^{b\text{-}t}$, *pnd*) (background strain used for injecting *3OHKT* targeting TALEN mRNA) are shown as control. (B) Sequences of small, induced insertions and deletions of G1 mutant females shown in (A). The inserted sequences are underlined.

A

| | sequence | Indel |
|---|---|---|
| Wild-type | TTTTTTCAGGGGTCCTCTACGAAGAAACAGGCAGCTACGTGTATTGTTTCTTGGCAATGAGTTTTT | none |
| G2 male #6 | TTTTTTCAGGGGTCCTCTACGAAGAAAC----------GTGTATTGTTTCTTGGCAATGAGTTTTT | Δ10 |
| G2 male #12 | TTTTTTCAGGGGTCCTCTACGAAGAAACA----GCTACGTGTATTGTTTCTTGGCAATGAGTTTTT | Δ4 |
|  | TTTTTTCAGGGGTCCTCTACGAAGAAAC----------GTGTATTGTTTCTTGGCAATGAGTTTTT | Δ10 |

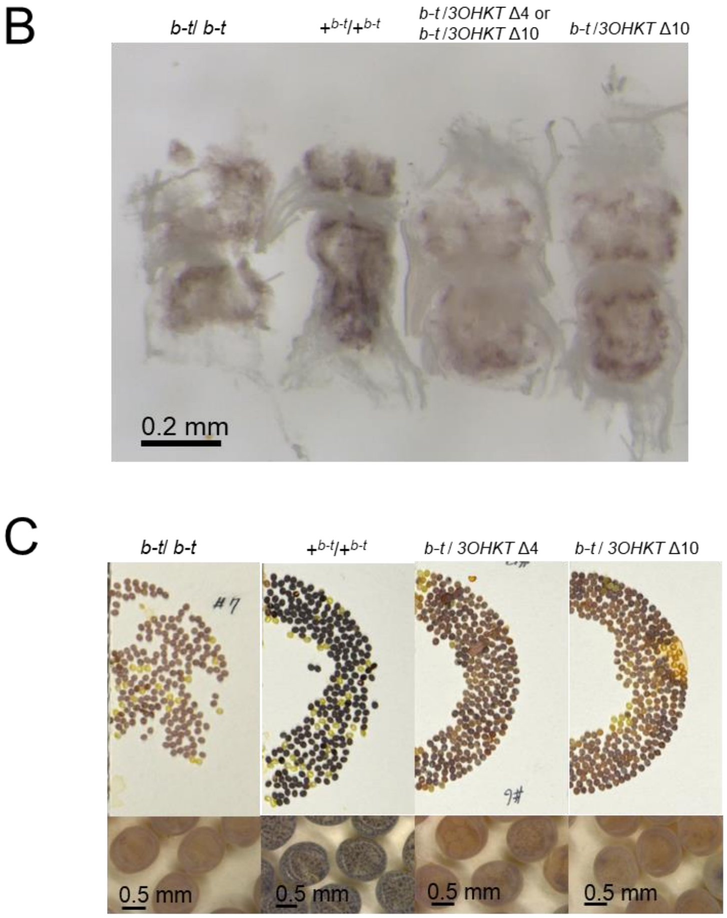


**Fig. S6. Complementation test of *b-t* mutant female with *KWMTBOMO07980* (*3OHKT*) deficient male.**

(A)Sequences of small, induced insertions and deletions in *3OHKT* for G2 males used in the complementation test. The inserted and deleted sequences are underlined and dotted line, respectively.

(B) The fifth-larval ganglion of *b-t* e30 (left), wild-type C108 ($+^{b\text{-}t}$, middle), and larvae derived from crosses between *b-t* (e30) and *3OHKT* knockout (Δ4/ Δ10, Δ10 homozygous) G2 individuals (right).

(C) Eggs laid by *b-t* (e30 strain, left), wild-type ($+^{b\text{-}t}$, *pnd*, middle), and females derived from crosses between *b-t* (e30) and *3OHKT* knockout (Δ4/ Δ10, Δ10 homozygous, genotype confirmed by genomic PCR and sequencing) G2 individuals (right).

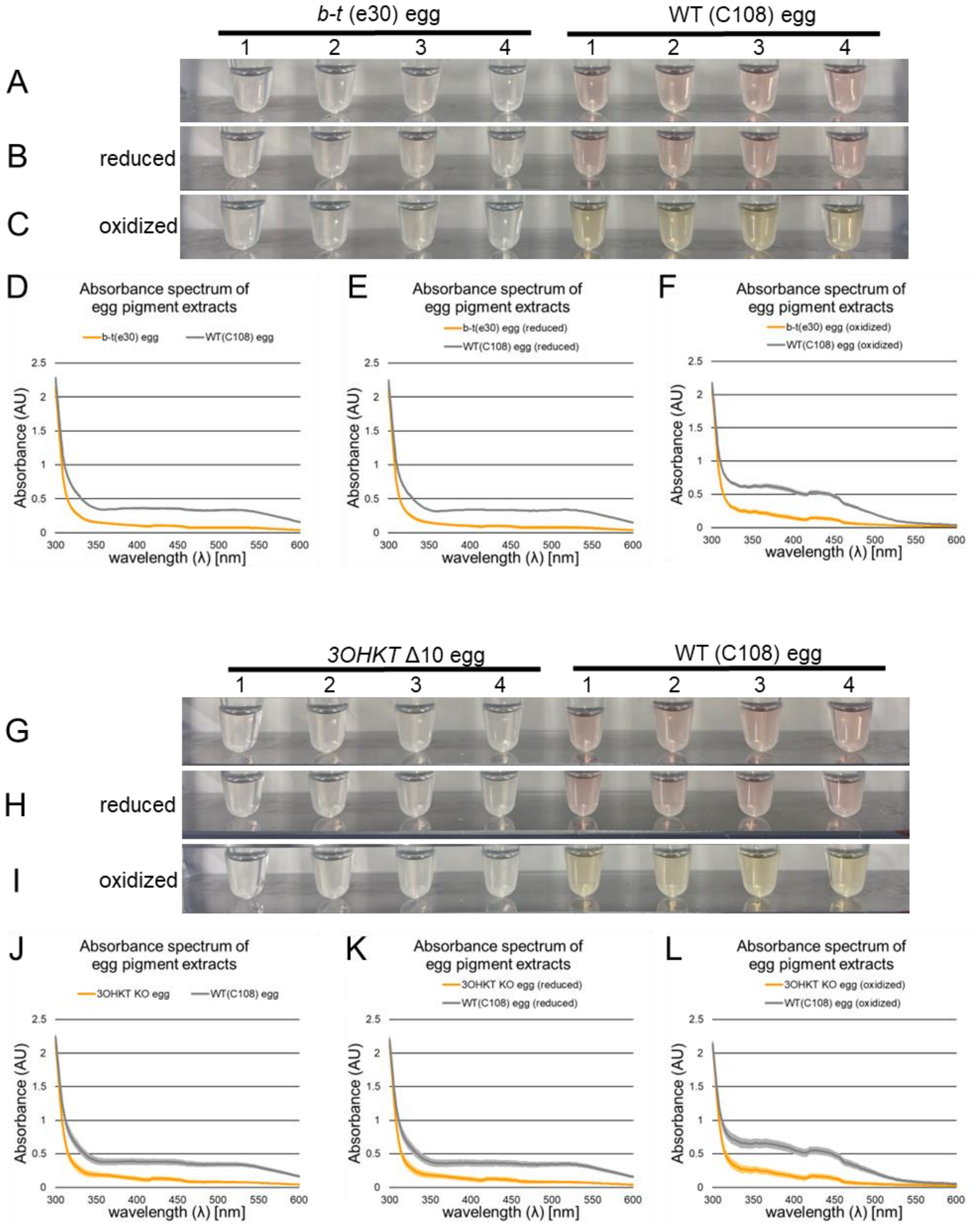


**Fig. S7. Ommochrome pigments in eggs of the wild-type, *b-t* mutant strain, and *3OHKT* knockout strain.**

C108 strain was used as wild-type control. The absorption spectrum of egg pigments in original extract (A, D, G, J), reduced state (B, E, H, K), and oxidized state (C, F, I, L). Group means and standard deviations are presented for the absorption spectrum of egg pigments (D-F, J-L). Four biological replicates were measured for each group. $p<0.05$ for Student's *t*-test implemented at 360, 380 450, 480 nm.

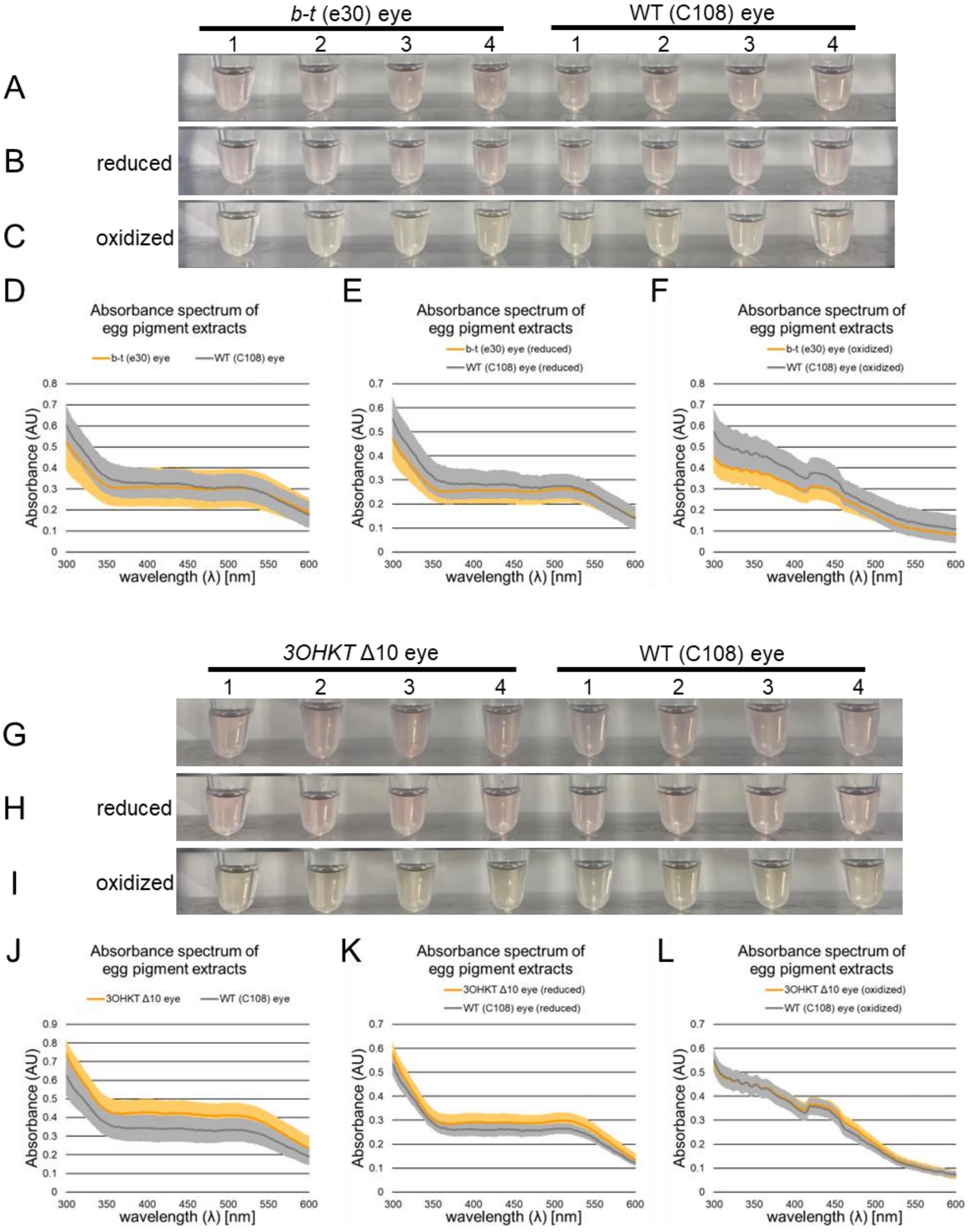


**Fig. S8. Ommochrome pigments in compound eyes of the wild-type, *b-t* mutant strain, and *3OHKT* knockout strain.**
p50T strain was used as wild-type control. The absorption spectrum of eye pigments in original extract (A, D, G, J), reduced state (B, E, H, K), and oxidized state (C, F, I, L). Group means and standard deviations are presented for the absorption spectrum of egg pigments (D-F, J-L). Four biological replicates were measured for each group. No significant differences detected by Student's t-test implemented at 360, 380 450, 480 nm between wild-type and *b-t* mutant strain or the *3OHKT* Δ10.

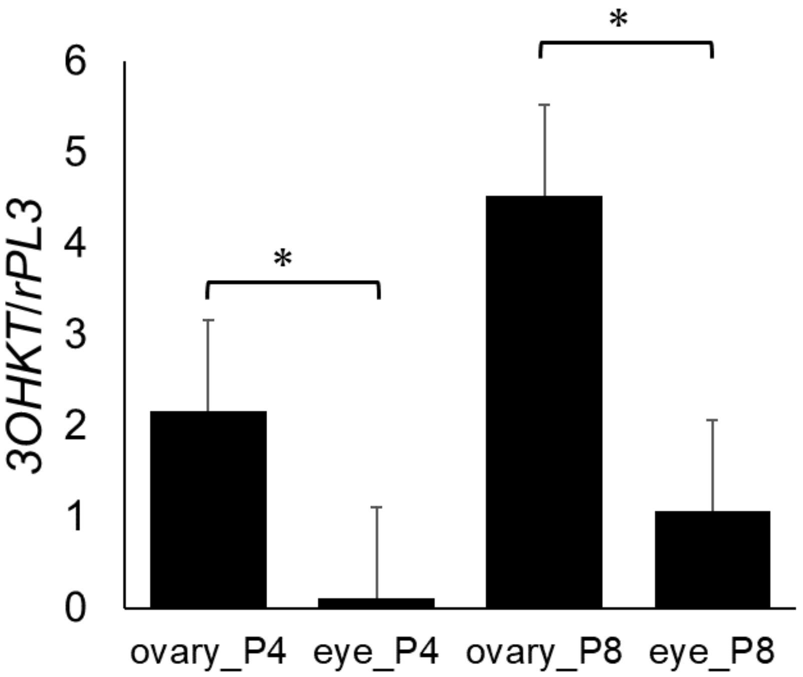


**Fig. S9. Expression level of *3OHKT* in pupal ovaries and eyes of *B. mori*.**
Relative expression profiles of *3OHKT* in the wild-type ovary and compound eye at pupa day 4 and 8. Expression of *ribosomal protein L3* (*rpL3*) was used as an internal control. Error bars indicate the SD (n=4). * $P < 0.05$ for Student's *t*-test.

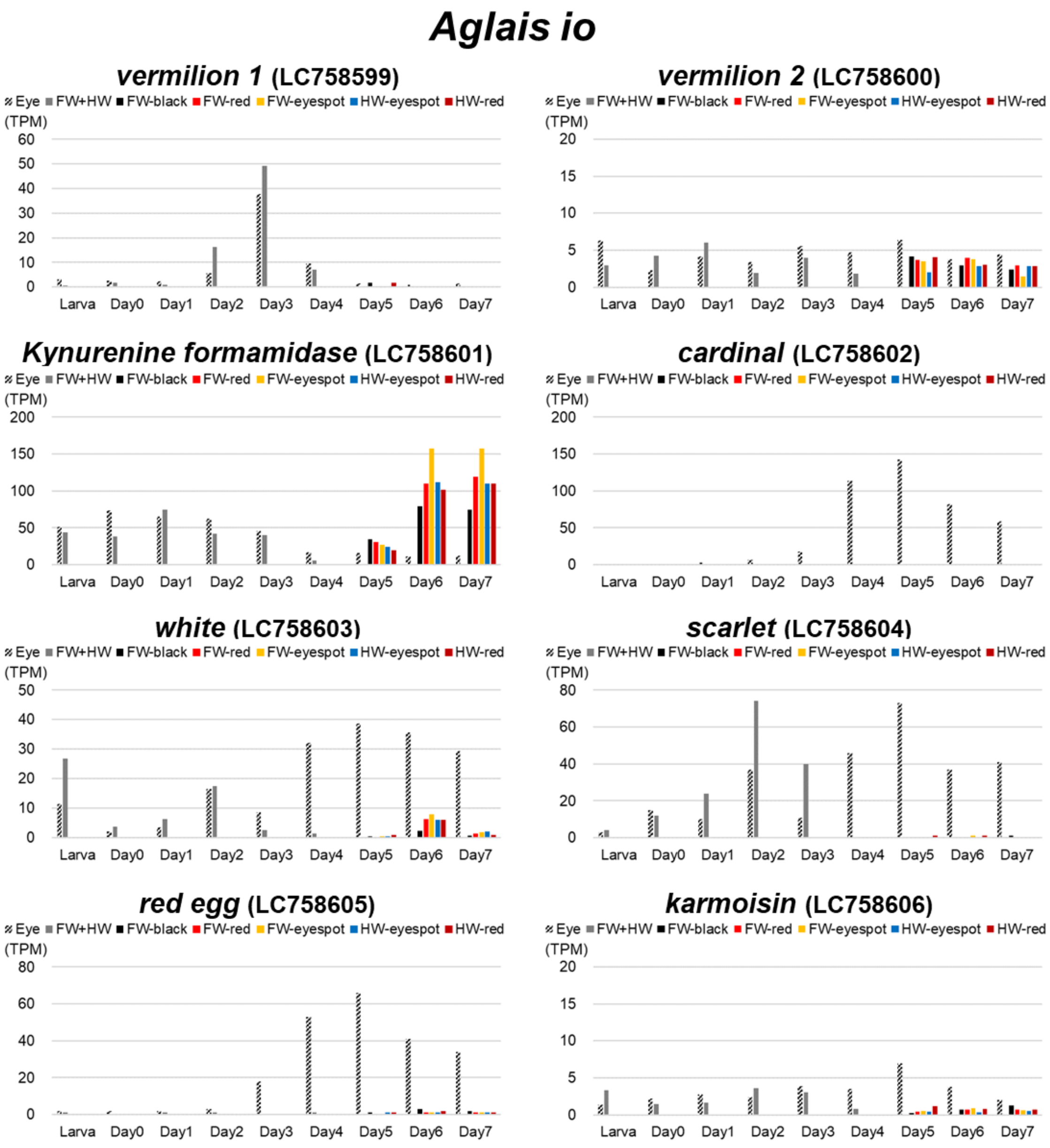


**Fig. S10 Expression profiles of ommochrome related genes in wings and eyes of the peacock butterfly *Aglais io*.**
For the region of the markings, refer to Fig.6A

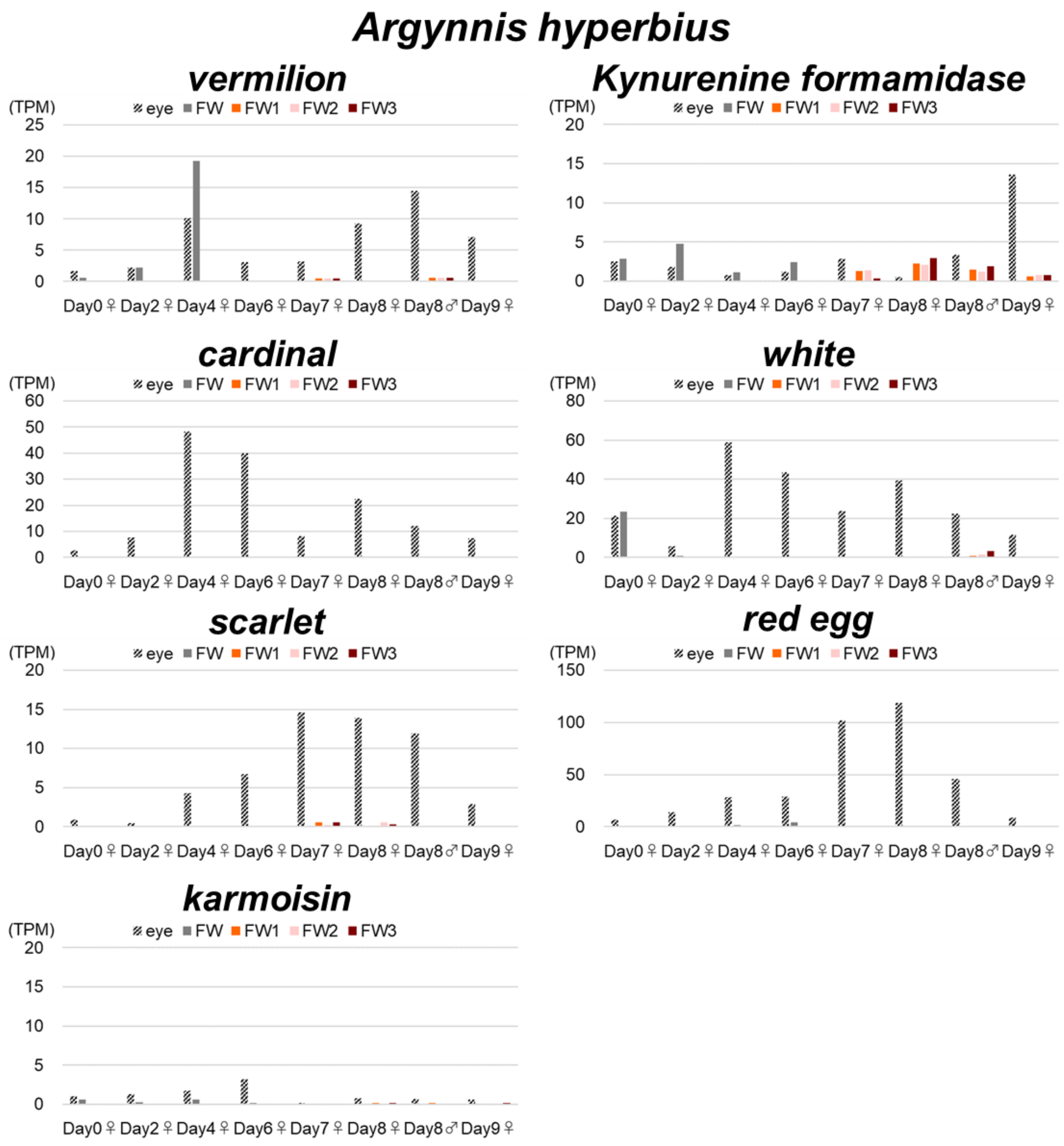


**Fig. S11 Expression profiles of ommochrome related genes in wings and eyes of the Indian fritillary butterfly *Argynnis hyperbius.***
For the region of the markings, refer to Fig.6C.

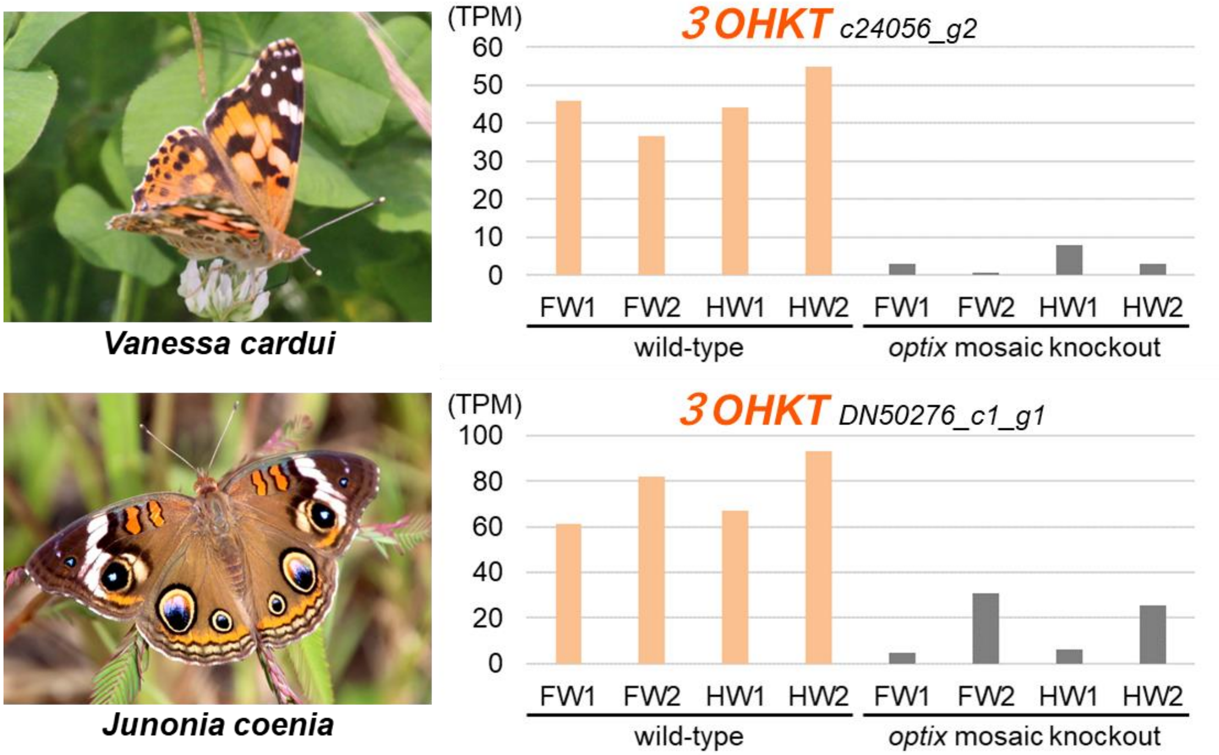


**Fig. S12 Expression levels of *3OHKT* gene in pupal wings of the wild-type and the *optix* mosaic knockout individuals in the painted lady butterfly *Vanessa cardui* and the buckeye butterfly *Junonia coenia.***
FW: forewing, HW: hindwing. RNA sequencing data are from Zhang et al. (2017).

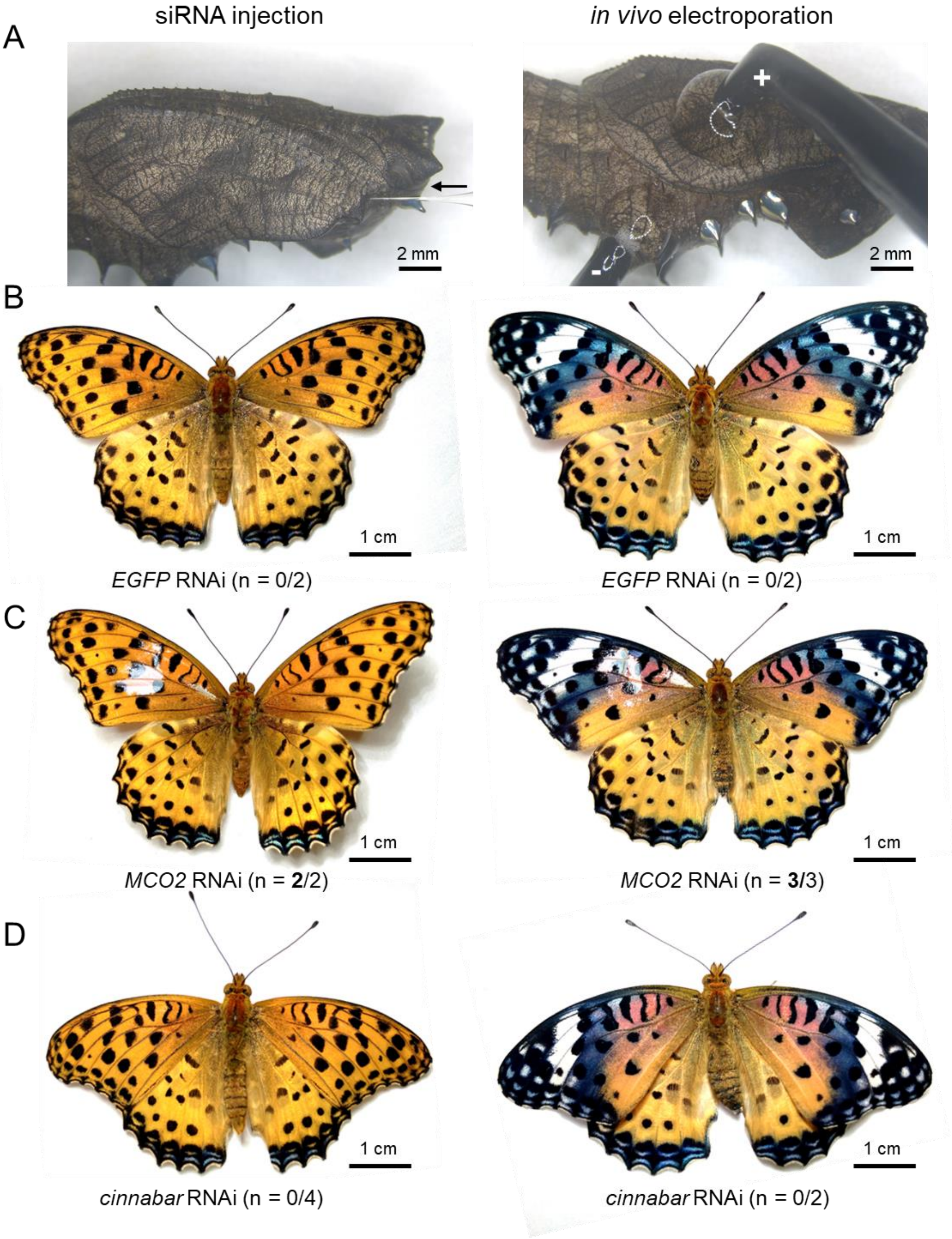


**Fig. S13. RNAi phenotypes on the wings of *A. hyperbius*.** (A) Electroporation-mediated RNAi method. After siRNA injection, electroporation is performed by negative (-) and positive (+) electrodes with droplets of ultrasound gel. (B) *EGFP* RNAi phenotype (negative control). (C) *MCO2* RNAi phenotype. (D) *cinnabar* RNAi phenotype. The Numbers in parentheses indicate (number of individuals affected by RNAi / number of emerged adults).

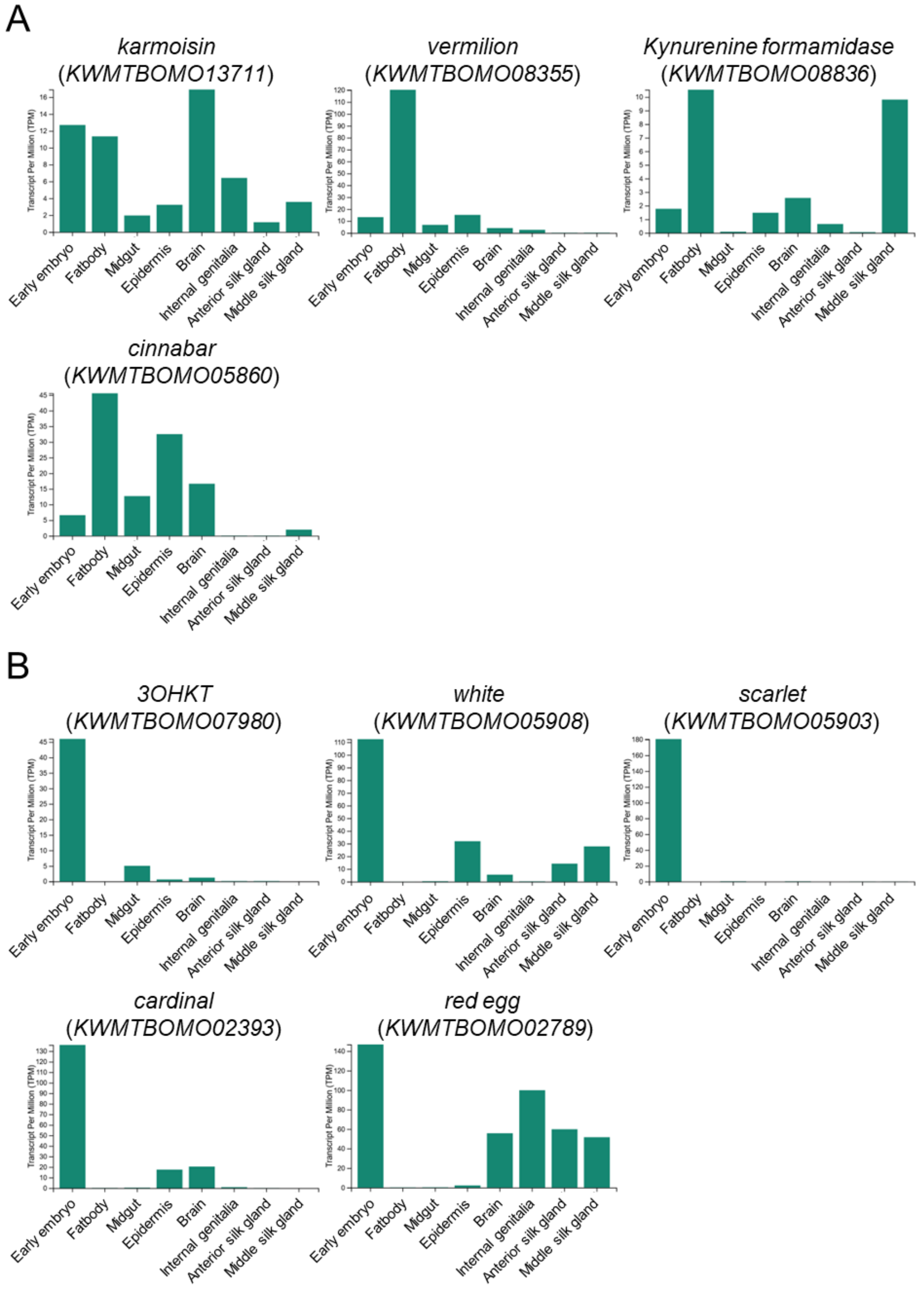


**Fig. S14. Expression data of ommochrome synthesis genes and transporters in *B. mori*.** Tissues were sampled at 5th instar larvae except for early embryo and Internal genitalia (adult). Data was obtained from silkbase (https://silkbase.ab.a.u-tokyo.ac.jp/cgi-bin/index.cgi). (A)Expression data of genes involved in 3OHK synthesis. (B) Expression data of genes involved in ommochrome pathway after 3OHK synthesis.

**Table S1.** ***A. hyperbius*** **siRNA list**

| | sense strand | antisense strand |
|---|---|---|
| *3OHKT* | GUG GUA UUG CUC UGA AAA AAU | UUU UUC AGA GCA AUA CCA CUG |
| *cinnabar* | GAA UUC UCG ACU ACU AGA UGG | AUC UAG UAG UCG AGA AUU CUU |
| *MCO2* | UUG CAU UUG GUG UAC AUA CUU | GUA UGU ACA CCA AAU GCA ACU |
| *EGFP* | GCA UCA AGG UGA ACU UCA AGA | UUG AAG UUC ACC UUG AUG CCG |

### Table S2. RNA sequencing sample list

| No. | Species | Sex | Developmental stage | Region | No. of pairs | Sequence | Accession No. |
|---|---|---|---|---|---|---|---|
| 1 | *Aglais io* | Male | Final instar larva | Head | 8,308,567 | HiSeq 150 bp paired | DRR443143 |
| 2 | *Aglais io* | Male | Final instar larva | Wing discs | 6,962,038 | HiSeq 150 bp paired | DRR443144 |
| 3 | *Aglais io* | Female | Pupa, day 0 | Compound eyes | 9,446,043 | HiSeq 150 bp paired | DRR443145 |
| 4 | *Aglais io* | Female | Pupa, day 0 | Wings | 8,679,410 | HiSeq 150 bp paired | DRR443146 |
| 5 | *Aglais io* | Female | Pupa, day 1 | Compound eyes | 8,928,685 | HiSeq 150 bp paired | DRR443147 |
| 6 | *Aglais io* | Female | Pupa, day 1 | Wings | 9,280,591 | HiSeq 150 bp paired | DRR443148 |
| 7 | *Aglais io* | Female | Pupa, day 2 | Compound eyes | 11,497,468 | HiSeq 150 bp paired | DRR443149 |
| 8 | *Aglais io* | Female | Pupa, day 2 | Wings | 12,829,534 | HiSeq 150 bp paired | DRR443150 |
| 9 | *Aglais io* | Female | Pupa, day 3 | Compound eyes | 8,282,633 | HiSeq 150 bp paired | DRR443151 |
| 10 | *Aglais io* | Female | Pupa, day 3 | Wings | 10,045,654 | HiSeq 150 bp paired | DRR443152 |
| 11 | *Aglais io* | Female | Pupa, day 4 | Compound eyes | 10,503,500 | HiSeq 150 bp paired | DRR443153 |
| 12 | *Aglais io* | Female | Pupa, day 4 | Wings | 14,311,679 | HiSeq 150 bp paired | DRR443154 |
| 13 | *Aglais io* | Female | Pupa, day 5 | Compound eyes | 9,195,236 | HiSeq 150 bp paired | DRR443155 |
| 14 | *Aglais io* | Female | Pupa, day 5 | Forewing region 1 | 9,596,259 | HiSeq 150 bp paired | DRR443156 |
| 15 | *Aglais io* | Female | Pupa, day 5 | Forewing region 2 | 10,624,779 | HiSeq 150 bp paired | DRR443157 |
| 16 | *Aglais io* | Female | Pupa, day 5 | Forewing region 3 | 10,717,606 | HiSeq 150 bp paired | DRR443158 |
| 17 | *Aglais io* | Female | Pupa, day 5 | Hindwing region 1 | 12,983,880 | HiSeq 150 bp paired | DRR443159 |
| 18 | *Aglais io* | Female | Pupa, day 5 | Hindwing region 2 | 7,224,016 | HiSeq 150 bp paired | DRR443160 |
| 19 | *Aglais io* | Female | Pupa, day 6 | Compound eyes | 10,795,131 | HiSeq 150 bp paired | DRR443161 |
| 20 | *Aglais io* | Female | Pupa, day 6 | Forewing region 1 | 10,877,459 | HiSeq 150 bp paired | DRR443162 |
| 21 | *Aglais io* | Female | Pupa, day 6 | Forewing region 2 | 10,203,040 | HiSeq 150 bp paired | DRR443163 |
| 22 | *Aglais io* | Female | Pupa, day 6 | Forewing region 3 | 6,276,676 | HiSeq 150 bp paired | DRR443164 |
| 23 | *Aglais io* | Female | Pupa, day 6 | Hindwing region 1 | 12,041,588 | HiSeq 150 bp paired | DRR443165 |
| 24 | *Aglais io* | Female | Pupa, day 6 | Hindwing region 2 | 8,963,887 | HiSeq 150 bp paired | DRR443166 |
| 25 | *Aglais io* | Male | Pupa, day 7 | Compound eyes | 8,090,285 | HiSeq 150 bp paired | DRR443167 |
| 26 | *Aglais io* | Male | Pupa, day 7 | Forewing region 1 | 8,421,640 | HiSeq 150 bp paired | DRR443168 |
| 27 | *Aglais io* | Male | Pupa, day 7 | Forewing region 2 | 9,178,614 | HiSeq 150 bp paired | DRR443169 |
| 28 | *Aglais io* | Male | Pupa, day 7 | Forewing region 3 | 13,359,905 | HiSeq 150 bp paired | DRR443170 |
| 29 | *Aglais io* | Male | Pupa, day 7 | Hindwing region 1 | 13,370,133 | HiSeq 150 bp paired | DRR443171 |
| 30 | *Aglais io* | Male | Pupa, day 7 | Hindwing region 2 | 8,844,660 | HiSeq 150 bp paired | DRR443172 |
| 31 | *Argynnis hyperbius* | Female | Pupa, day 0 | Compound eyes | 9,312,268 | HiSeq 150 bp paired | DRR443173 |
| 32 | *Argynnis hyperbius* | Female | Pupa, day 0 | Forewing | 9,221,825 | HiSeq 150 bp paired | DRR443174 |
| 33 | *Argynnis hyperbius* | Female | Pupa, day 2 | Compound eyes | 12,244,061 | HiSeq 150 bp paired | DRR443175 |
| 34 | *Argynnis hyperbius* | Female | Pupa, day 2 | Forewing | 9,694,796 | HiSeq 150 bp paired | DRR443176 |
| 35 | *Argynnis hyperbius* | Female | Pupa, day 4 | Compound eyes | 6,387,745 | HiSeq 150 bp paired | DRR443177 |
| 36 | *Argynnis hyperbius* | Female | Pupa, day 4 | Forewing | 9,891,875 | HiSeq 150 bp paired | DRR443178 |
| 37 | *Argynnis hyperbius* | Female | Pupa, day 6 | Compound eyes | 4,592,237 | HiSeq 150 bp paired | DRR443179 |
| 38 | *Argynnis hyperbius* | Female | Pupa, day 6 | Forewing | 3,693,270 | HiSeq 150 bp paired | DRR443180 |
| 39 | *Argynnis hyperbius* | Female | Pupa, day 7 | Compound eyes | 11,634,694 | HiSeq 150 bp paired | DRR443181 |
| 40 | *Argynnis hyperbius* | Female | Pupa, day 7 | Forewing region 1 | 10,711,991 | HiSeq 150 bp paired | DRR443182 |
| 41 | *Argynnis hyperbius* | Female | Pupa, day 7 | Forewing region 2 | 10,832,168 | HiSeq 150 bp paired | DRR443183 |
| 42 | *Argynnis hyperbius* | Female | Pupa, day 7 | Forewing region 3 | 8,616,617 | HiSeq 150 bp paired | DRR443184 |
| 43 | *Argynnis hyperbius* | Female | Pupa, day 8 | Compound eyes | 9,806,157 | HiSeq 150 bp paired | DRR443185 |
| 44 | *Argynnis hyperbius* | Female | Pupa, day 8 | Forewing region 1 | 9,605,319 | HiSeq 150 bp paired | DRR443186 |
| 45 | *Argynnis hyperbius* | Female | Pupa, day 8 | Forewing region 2 | 11,674,784 | HiSeq 150 bp paired | DRR443187 |
| 46 | *Argynnis hyperbius* | Female | Pupa, day 8 | Forewing region 3 | 9,502,487 | HiSeq 150 bp paired | DRR443188 |
| 47 | *Argynnis hyperbius* | Male | Pupa, day 8 | Compound eyes | 9,250,450 | HiSeq 150 bp paired | DRR443189 |
| 48 | *Argynnis hyperbius* | Male | Pupa, day 8 | Forewing region 1 | 8,721,119 | HiSeq 150 bp paired | DRR443190 |
| 49 | *Argynnis hyperbius* | Male | Pupa, day 8 | Forewing region 2 | 6,998,194 | HiSeq 150 bp paired | DRR443191 |
| 50 | *Argynnis hyperbius* | Male | Pupa, day 8 | Forewing region 3 | 8,695,813 | HiSeq 150 bp paired | DRR443192 |
| 51 | *Argynnis hyperbius* | Female | Pupa, day 9 | Compound eyes | 13,863,519 | HiSeq 150 bp paired | DRR443193 |
| 52 | *Argynnis hyperbius* | Female | Pupa, day 9 | Forewing region 1 | 10,151,650 | HiSeq 150 bp paired | DRR443194 |
| 53 | *Argynnis hyperbius* | Female | Pupa, day 9 | Forewing region 2 | 13,378,251 | HiSeq 150 bp paired | DRR443195 |
| 54 | *Argynnis hyperbius* | Female | Pupa, day 9 | Forewing region 3 | 11,597,151 | HiSeq 150 bp paired | DRR443196 |
| 55 | *Bombyx mori* | Mixed | 0 hour after egg laying | *b-t* strain, whole embryo | 6,295,336 | HiSeq 100 bp paired | DRR277731 |
| 56 | *Bombyx mori* | Mixed | 24 hour after egg laying | *b-t* strain, whole embryo | 6,102,964 | HiSeq 100 bp paired | DRR277732 |
| 57 | *Bombyx mori* | Mixed | 48 hour after egg laying | *b-t* strain, whole embryo | 6,869,139 | HiSeq 100 bp paired | DRR277733 |
| 58 | *Bombyx mori* | Mixed | 72 hour after egg laying | *b-t* strain, whole embryo | 5,500,803 | HiSeq 100 bp paired | DRR277734 |
| 59 | *Bombyx mori* | Mixed | 0 hour after egg laying | *p50T* strain, whole embryo | 15,809,762 | HiSeq 100 bp paired | DRR030435 |
| 60 | *Bombyx mori* | Mixed | 24 hour after egg laying | *p50T* strain, whole embryo | 17,905,594 | HiSeq 100 bp paired | DRR030436 |
| 61 | *Bombyx mori* | Mixed | 48 hour after egg laying | *p50T* strain, whole embryo | 18,897,878 | HiSeq 100 bp paired | DRR030437 |
| 62 | *Bombyx mori* | Mixed | 72 hour after egg laying | *p50T* strain, whole embryo | 18,575,189 | HiSeq 100 bp paired | DRR030438 |
| 63 | *Bombyx mori* | Mixed | 0 hour after egg laying | *C108* strain, whole embryo | 21,841,083 | HiSeq 100 bp paired | DRR030431 |
| 64 | *Bombyx mori* | Mixed | 24 hour after egg laying | *C108* strain, whole embryo | 18,503,807 | HiSeq 100 bp paired | DRR030432 |
| 65 | *Bombyx mori* | Mixed | 48 hour after egg laying | *C108* strain, whole embryo | 21,686,965 | HiSeq 100 bp paired | DRR030433 |
| 66 | *Bombyx mori* | Mixed | 72 hour after egg laying | *C108* strain, whole embryo | 16,123,187 | HiSeq 100 bp paired | DRR030434 |

**Table S3. Primer list**

| primer usage/name | sequence |
|---|---|
| **genotyping** | |
| **Bmb-t_wt-genotyping-F** | ACGACTCCATTAGGGAAATT |
| **Bmb-t_wt-genotyping-R** | ACGATTTGTACGATAGACAA |
| **T7** | TAATACGACTCACTATAGGG |
| **SP6-2** | AGGTGACACTATAGAATACTC |
| **Plasmid construction** | |
| **pBac-XbaI-3xflag-b-t_wt-F-Gibson** | AGGATGACGATGACAAGAGTATGGATGAGGCTTTCGCG |
| **pBac-XbaI-3xflag-b-t_wt-R-Gibson** | ACCTTCGAACCGCGGGCCCTTTAATTAGAATTACGATTTGTACGATAGACAAAATAC |
| **pBac-XbaI_C_3xflag-b-t_wt-F-Gibson** | ACAGTGGCGGCCGCTCGAGTATGGATGAGGCTTTCGCG |
| **pBac-XbaI_C_3xflag-b-t_wt-R-Gibson** | ACCTTCGAACCGCGGGCCCTGATTAGAATTACGATTTGTACGATAGACAAAATAC |
| **pBac-b-t_wt-seq1** | TTCGAATTTAAAGCTTGGTACC |
| **pBac-b-t_wt-seq2** | TAATTAGAGGAGCCGGCTTTG |
| **pBac-b-t_wt-seq3** | TGGGTTTCGCTTTCGTTTC |
| **pBac-b-t_wt-seq4** | TACGCGTAGAATCGAGACCG |
| **EBV_rev_primer** | GTGGTTTGTCCAAACTCATC |
| **OpIE2 reverse** | GACAATACAAACTAAGATTTAGTCAG |
| **genomic PCR** | |
| **b-t_intron-F7** | AGTAGGACATCTCATCGCATC |
| **b-t_exon9-R2** | GTAACAAACAAAAGCACTGTCAC |
| **RT-PCR** | |
| **Bmb-t_wt-exon8-F** | TATGGCAGCCCCTGATA |
| **Bmb-t_wt-exon9-R** | AAAAACTCATTGCCAAGAAAC |
| **rpL3-3** | TGCGTCCAAGCTCATCCTGC |
| **rpL3-5** | AGCACCCCGTCATGGGTCTA |
| **real-time PCR** | |
| **Bmb-t_wt-exon8-F** | TATGGCAGCCCCTGATA |
| **Bmb-t_wt-exon9-R** | AAAAACTCATTGCCAAGAAAC |
| **qPCR-rpL3-F** | ATCAAGGGTTGCTGCATGGGACCT |
| **qPCR-rpL3-R** | GGTTGATCTTTTCTAGTGCAGCCCTC |
| **Heteroduplex analysis** | |
| **Bmb-t_wt-exon9-F** | TTTTTTCAGGGGTCCTCTAC |
| **Bmb-t_wt-longPCR-R** | ATTACCAAAACACTTATTGGT |

### SI References


1. H. Mon, *et al.*, Effective RNA interference in cultured silkworm cells mediated by overexpression of Caenorhabditis elegans SID-1. *RNA Biol.* **9**, 40–6 (2012).
2. G. Okude, *et al.*, Molecular mechanisms underlying metamorphos is in the most-ancestral winged insect. *Proc. Natl. Acad. Sci. U. S. A.*, **119**, e2114773119 (2022).
3. J. T. Robinson *et al.*, Integrative Genomics Viewer. *Nat Biotechnol.,* **29**, 24–26 (2011).
4. M. G. Grabherr et al., Full-length transcriptome assembly from RNA-Seq data without a reference genome. *Nat Biotechnol.,* **29**, 644–652 (2011).
5. M. Kawamoto, et al., High-quality genome assembly of the silkworm, Bombyx mori. *Insect Biochem. Mol. Biol.* **107**, 53–62 (2019)
6. R. Patro, G. Duggal, M. I. Love, R. A. Irizarry, C. Kingsford, Salmon provides fast and bias-aware quantification of transcript expression. Nat Methods 14, 417–419 (2017).
7. L. Käll, A. Krogh, E. L. L. Sonnhammer, Advantages of combined transmembrane topology and signal peptide prediction--the Phobius web server. *Nucleic Acids Res.*, **35**, W429-32 (2007).
8. Y. Takasu, *et al.*, Efficient TALEN Construction for Bombyx mori Gene Targeting. *PLoS One* **8**, 1–11 (2013).
9. T. Cermak, *et al.*, Efficient design and assembly of custom TALEN and other TAL effector-based constructs for DNA targeting. *Nucleic Acids Res.* **39**, e82 (2011).
10. K. Tamura, G. Stecher, D. Peterson, A. Filipski, S. Kumar, MEGA6: Molecular Evolutionary Genetics Analysis version 6.0. *Mol. Biol. Evol.* **30**, 2725–9 (2013).
11. S. Capella-Gutiérrez, J. M. Silla-Martínez, T. Gabaldón, trimAl: a tool for automated alignment trimming in large-scale phylogenetic analyses. *Bioinformatics* **25**, 1972–3 (2009).
12. A. Stamatakis, RAxML version 8: a tool for phylogenetic analysis and post-analysis of large phylogenies. *Bioinformatics* **30**, 1312–3 (2014).
13. A. S. Tanabe, Kakusan4 and Aminosan: two programs for comparing nonpartitioned, proportional and separate models for combined molecular phylogenetic analyses of multilocus sequence data. *Mol. Ecol. Resour.* **11**, 914–21 (2011).
